# Structural basis of apoptosis induction by the mitochondrial voltage dependent anion channel

**DOI:** 10.1101/2025.04.15.648943

**Authors:** Melina Daniilidis, Umut Günsel, Georgios Broutzakis, Robert Janowski, Kira D. Leitl, Kai Fredriksson, Dierk Niessing, Christos Gatsogiannis, Franz Hagn

## Abstract

The voltage-dependent anion channel (VDAC) is the main gateway for metabolites across the mitochondrial outer membrane^1^. In addition, VDAC oligomers have been associated with apoptosis at mitochondrial stress conditions^2^. However, the mechanistic and structural basis of VDAC’s capability to induce apoptosis pathways remains poorly understood. Here, we show with biochemical and structural methods that VDAC1 oligomerization triggers the dissociation of its N-terminal α-helix (VDAC1-N) from the channel interior. We used advanced lipid nanodiscs as a tool to selectively trap VDAC1 in its canonical helix-inserted and helix-exposed state to facilitate a structural characterization of both conformations by cryo-electron microscopy. The results show that slight changes in the shape and dynamics of the VDAC1 β-barrel suffice to release the N-terminal helix to the channel exterior. This conformational switch addresses the long-standing question how VDAC1 can regulate partner protein binding. To confirm this hypothesis, we performed interaction studies between VDAC1 in both conformational states and the anti-apoptotic partner protein BclxL using nuclear magnetic resonance spectroscopy and could detect binding only for the helix-exposed state. These insights enabled the X-ray structure determination of the BclxL-VDAC1-N complex at high resolution and provided atomistic details on the VDAC1-N binding mode at the BH3-groove in BclxL. Further biochemical assays showed that VDAC1-N promotes pore formation of the pro-apoptotic Bcl2 protein Bak by neutralizing BclxL’s inhibitory activity. These findings suggest that stress-induced oligomerization of VDAC can trigger the exposure of its N-terminal α-helix leading to the neutralization of anti-apoptotic Bcl2 proteins. This mode-of-action is reminiscent of BH3-only sensitizer Bcl2 proteins^3^ that are efficient inducers of Bax/Bak-mediated mitochondrial outer membrane permeabilization and ultimately apoptosis.

## Introduction

The mitochondrial voltage-dependent anion channel (VDAC) is the major transit pore of (energy-) metabolites and metal ions across the outer mitochondrial membrane^1^. VDAC plays a major role in the molecular pathology of e.g. Alzheimer’s^4^ and Parkinson’s disease or cancer^5^, and cardiac ischemia–reperfusion injury^6,7^. VDAC1 is a porin composed of a 19-strand β-barrel and an N-terminal α-helix (VDAC1-N) attached to the inside of the pore^8-10^. The inner surface of the β-barrel is decorated with positive charges that provide a preference of the channel for negatively charged substrates, such as glutamate^11^ and ATP^12^, while also transporting positively charged metabolites like acetylcholine or dopamine^11^. Electrophysiology experiments showed that VDAC1 can adopt an open state at low to zero and a closed state at high membrane potential^13^, shifting the channel’s selectivity to small cations^14-16^.

In addition to its role in mitochondrial metabolite transport, VDAC1 has also been implicated as a key player in programmed cell death. VDAC1 has been reported to act as an inducer of apoptosis^2^ via mitochondrial outer membrane permeabilization (MOMP)^17^, a process that enables the release of pro-apoptotic proteins from the intermembrane space (IMS) to activate caspases^18^. However, the inner pore of the VDAC1 monomer has only a diameter of 1.5 to ∼3.0 nm^8-10,19,20^, which renders it too small for the release of pro-apoptotic proteins.

Mitochondrial damage caused by various inducers of apoptosis, including inhibitors of the respiratory chain or hydrogen peroxide, have been reported to cause VDAC1 oligomerization and eventually apoptosis^2^. Thus, it has been suggested that such oligomers might form large-enough pores in the membrane to execute MOMP^21^ and even release mitochondrial DNA fragments^22^. VDAC1 has a tendency to form dimers and oligomers in detergent solution^23^ and native membranes^20^ but stable oligomerization in a cellular environment requires additional stimuli such as altered mitochondrial lipid composition^24,25^, increased Ca^2+^ levels^26,27^, low pH^28^ or oxidative stress^22,29^.

The interaction between VDAC1 and the anti-apoptotic Bcl2 protein BclxL has been reported to be considerably enhanced by apoptotic stimuli^30,31^, which was confirmed by *in vitro* ^23^ and cellular studies^32,33^. This interaction was reported to be essential for the execution of apoptosis under mitochondrial stress^34^. The Bcl2 protein family is the canonical system to induce MOMP^3,35^, where the activation of the pore-forming Bcl2 proteins Bak and Bax leads to large membrane pores of 40 nm to 1 µm in diameter^36,37^ that allow for the exit of folded proteins or even mitochondrial DNA^38^. This process is constantly inhibited by the anti-apoptotic Bcl2 protein members, such as BclxL, Bcl2 or Mcl1^3^. Thus, a protein that can neutralize the anti-apoptotic Bcl2 members, inevitably leads to apoptosis via MOMP induced by activated Bax/Bak^39^.

Despite this large body of evidence, the exact mechanistic role of VDAC in the intrinsic induction pathways of apoptosis remains highly controversial^17^. Structural studies of the complex between VDAC1 and BclxL are essential to reveal the mechanistic details of a possible connection between VDAC and the Bcl2 protein family in the regulation of apoptosis.

Here, we used structural and biochemical methods to explore the mechanistic basis of MOMP induction by VDAC1 via its interaction with the anti-apoptotic Bcl2 protein BclxL. We show that VDAC1 oligomerization in negatively charged detergents or lipids leads to the exposure of the VDAC1 N-terminal α-helix (VDAC1-N). Using cryo-EM in circularized lipid nanodiscs of different sizes^40,41^, we were able to structurally characterize VDAC1 in different conformational states where VDAC1-N is either bound inside the pore or exposed to the pore exterior. Using NMR, we also demonstrated that binding to BclxL is only possible if VDAC1-N becomes exposed. This observation was further confirmed by binding assays and a high-resolution crystal structure of the VDAC1-N – BclxL complex. Liposome pore-forming assays provided evidence that VDAC1-N can dissociate the inhibitory complex between BclxL and the pore-forming Bcl2 protein Bak, restoring its pore forming activity. We conclude that apoptotic stimuli that induce VDAC1 oligomerization lead to the exposure of VDAC1-N, which acts as a sensitizer BH3 protein to inhibit anti-apoptotic Bcl2 proteins. As a result, VDAC1-dependent MOMP can be executed by pro-apoptotic effector Bcl2 proteins.

## Results

### VDAC1 oligomerization leads to exposure of its N-terminal α-helix

First, we wondered whether the N-terminal α-helix of VDAC1, the putative interaction site with Bcl2 proteins^33^, can become exposed under apoptotic conditions, i.e. in the oligomeric state of VDAC1. VDAC1 oligomerization has been shown to be a marker for apoptosis^2^ (**Fig. 1a**). In the previously published structures of VDAC1 the N-terminal α-helix is stably attached to the pore interior^8-10^. Thus, a marked conformational change must take place to enable the helix to dissociate from the β-barrel wall. To address this question, we first used chemical crosslinking experiments to identify conditions that favor VDAC1 oligomerization in detergent micelles and liposomes. We screened several detergents for promoting VDAC1 oligomerization which we probed by chemical crosslinking experiments with the amino-selective crosslinker bissulfosuccinimidyl suberate (BS^3^) (**Fig. 1b**). The zwitterionic detergent LDAO that has been used for structural studies on VDAC1 and the milder detergent TritonX-100 both induce very moderate VDAC1 oligomerization. In the negatively charged bile acid cholate, VDAC1 oligomerization is strongly favored which contrasts the very similar zwitterionic bile acid derivate CHAPS. We show that in a liposome environment, the formation of very large VDAC1 oligomers can be facilitated by negatively charged lipids such as POPG (**Fig. S1c**). To probe whether VDAC1 oligomerization might be a trigger for the exposure of its N-terminal α-helix, we used the cysteine-specific maleimide-PEG40 (PM40) reagent and a cysteine-free VDAC1 variant where a single cysteine was re-introduced at position 6 (T6C). The latter is located at the N-terminal end of the α-helix that is not exposed to the pore exterior (**Fig. 1c**). Due to a calculated hydrodynamic radius of ∼6.2 nm^42^, the 40 kDa PEG polymer cannot access the cysteine inside the pore (see structural model in **Fig. 1c**) having an inner diameter of 1.5-3 nm. Thus, the modification reaction can only happen if the N-terminal part of VDAC1 becomes exposed to the solvent. In very good agreement with the oligomerization assay, PM40 modification of VDAC1 is very pronounced in the detergent cholate and to a much lesser extent in the other detergents that do not bear a net negative charge (**Fig. 1c**). This data suggests that VDAC1 oligomerization and the exposure of the N-terminal helix are directly connected, which would imply that in turn VDAC1 stabilization can possibly reduce its degree of oligomerization. To test this hypothesis, we used the well-characterized VDAC1-E73V variant and observed higher thermal stability than for wt-VDAC1 (**Fig. S2a**). VDAC1-E73V exhibits reduced dynamics in the β-barrel and a more stable attachment of the N-terminal α-helix^8,43^, facilitating its NMR structure determination^44,45^. In line with these previous data, this mutation caused a marked reduction in VDAC1 oligomerization in our experiments (**Fig. S1c**) and a lower degree of exposure of its N-terminal helix (**Fig. S1a**). If the bulky hydrophobic side chain of Leu10 within the N-terminal helix that mediates the interaction with the β-barrel was replaced by alanine (L10A) in VDAC1-E73V, helix exposure and degree of oligomerization were again comparable to wt-VDAC1 (**Fig. S1b**,**c**).

**Fig. 1.**
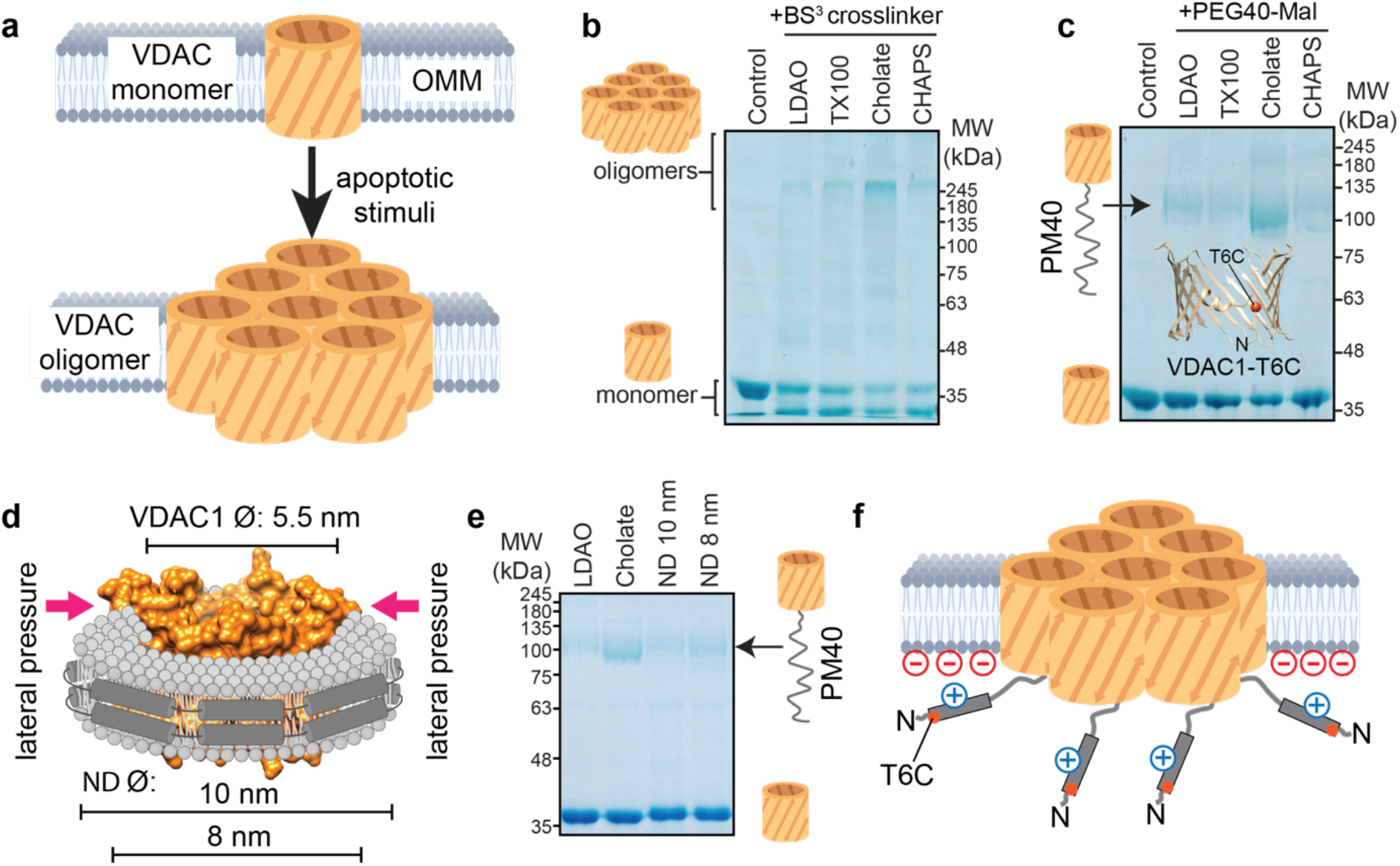
VDAC1 exposes its N-terminal α-helix in the oligomeric state. (a) Upon various apoptotic stimuli, VDAC1 was shown to form oligomers in the outer mitochondrial membrane (OMM). (b) VDAC1 oligomerization in detergent micelles as detected by amino-specific crosslinking with bissulfosuccinimidyl suberate (BS^3^) is favored by a negative net charge, as present in the detergent cholate. (c) Modification of VDAC1 at the sulfhydryl sidechain of cysteine 6 (T6C) in a cysteine-free background by maleimide-polyethylene glycol of 40 kDa (PM40) only occurs in the oligomeric form. (d)Lipid nanodiscs (ND) of different sizes mimic the oligomeric state of VDAC1 in various extents. VDAC1 is shown in orange, the membrane scaffold protein (MSP) in dark grey and lipids in light grey. (e)The smaller nanodisc (8 nm diameter) leads to a more pronounced exposure of cysteine 6 in VDAC1 than the 10 nm nanodisc, as probed by PM40 modification. (f) The PM40 data suggests that in the oligomeric state or if inserted into small lipid nanodiscs, VDAC1 exposes its N-terminal α-helix to the solvent. A negative net charge in the lipid headgroups facilitates helix exposure by electrostatic interactions with the α-helix.

To enable structural studies of VDAC1 in the helix inserted or exposed states, we explored the use of lipid nanodiscs. We used lipid nanodiscs of 10 and 8 nm diameter^46-48^ where VDAC1 with an outer diameter of ∼5.5 nm can be inserted together with a varying amount of residual lipids between the membrane protein and the membrane scaffold protein (MSP) (**Fig. 1d**). Thus, we anticipated that the structural constraint imposed by the nanodisc might induce a structural transformation as seen in detergent micelles without the necessity to adopt an oligomeric state. To detect helix exposure in VDAC1 in lipid nanodiscs we used the PM40 assay described above. This data shows that the exposed N-terminal helix conformation of VDAC1 can be stabilized in smaller nanodiscs as seen by the stronger band in the SDS-PAGE at ∼100kDa, representing the VDAC1-T6C-PM40 conjugate (**Fig. 1e, Fig. S1a**,**b**). Since VDAC1 is present mainly as a monomer in nanodiscs, the E73V mutation did not reduce helix exposure. Thus, the 8 nm nanodisc seems suitable to stabilize a helix-exposed VDAC1 structural state for cryo-EM and NMR studies.

In pure lipids, VDAC1 generally showed a rather low tendency to form oligomers, partially because higher amounts of VDAC1 could not be inserted into liposomes (max. ∼10 µM). Despite this limitation, we could detect the formation of very large oligomeric species prominently in the presence of the negatively charged lipid 1-palmitoyl-2-oleyl-glycero-3-phospho-glycerol (POPG), which also induced a higher exposure of the N-terminal helix as monitored by the PEG40 assay (**Fig. S1d, e**). The effect of negatively charged lipids can be rationalized by the positive net charge of VDAC1-N, suggesting that the helix, once exposed, can interact electrostatically with the membrane surface, as reported previously^49-51^. Thermal melting experiments with VDAC1 in nanodiscs of different sizes, where the α-helix is either located inside the pore (10 nm, MSP1D1) or exposed (8 nm, MSP1D1ΔH5), show that the exposed structural state is less stable (**Fig. S2b**), suggesting that the absence of the N-terminal helix at the barrel wall leads to structural instability of VDAC1 as previously monitored by a loss in functionality and voltage gating activity^34,52^. Our data indicate that VDAC1 oligomerization is the trigger for the dissociation of the N-terminal α-helix from the pore interior (**Fig. 1f**), whereas in the monomeric state, the helix is stably attached to the β-barrel.

### Cryo-EM structures of VDAC1 in lipid nanodiscs reveal conformational switching

Motivated by the biochemical experiments where the extent of α-helix exposure in 10 nm and 8 nm nanodiscs differed significantly, we further set out to characterize the structural states of VDAC1 using cryo-EM. For cryo-EM we used advanced lipid nanodiscs where the membrane scaffold protein is covalently circularized (cMSP), giving rise to enhanced stability and homogeneity but with a slightly larger (∼1 nm) diameter ^40,41,53^.

Reconstitution of VDAC1 in cMSP1D1 nanodiscs (11 nm diameter) followed by size exclusion chromatography (SEC) yielded two major peaks, both of which were analyzed using single-particle cryo-EM (**Fig. S3a**,**b**). The higher molecular weight peak eluting earlier in the SEC run corresponds to the monomeric state of VDAC1 (**Fig. 2a-d, Fig. S3**). For this species, we obtained a map at a resolution of 7 Å from 31,602 particles, showing the VDAC1 protomer residing at the lateral edge of the nanodisc (**Fig. S4, upper panel**). The N-terminal helix is resolved and located inside the pore at the expected position (**Fig. 2 c,d**). The monodisperse cryo-EM sample of the late-eluting peak corresponds to VDAC1 dimers (**Fig. 2e-h, Fig. S3e**). We were also able to obtain a map of the VDAC1 dimer at a resolution of 7.2 Å from 169,625 particles, with no symmetry imposed. The N-terminal helix of each protomer is resolved as a clear rod-like density localized within the interior of the barrel (**Figs. 2g, h**). Thereby, the nanodisc adopts an oval shape to accommodate the VDAC1 dimer, displaying direct interactions between the membrane scaffold protein and the two VDAC1 barrels. The dimerization interface of the two VDAC1 protomers is formed by the face of the β-barrel where the N-terminal helix is attached to, as seen previously in a crystal structure of human VDAC1^54^.

**Fig. 2.**
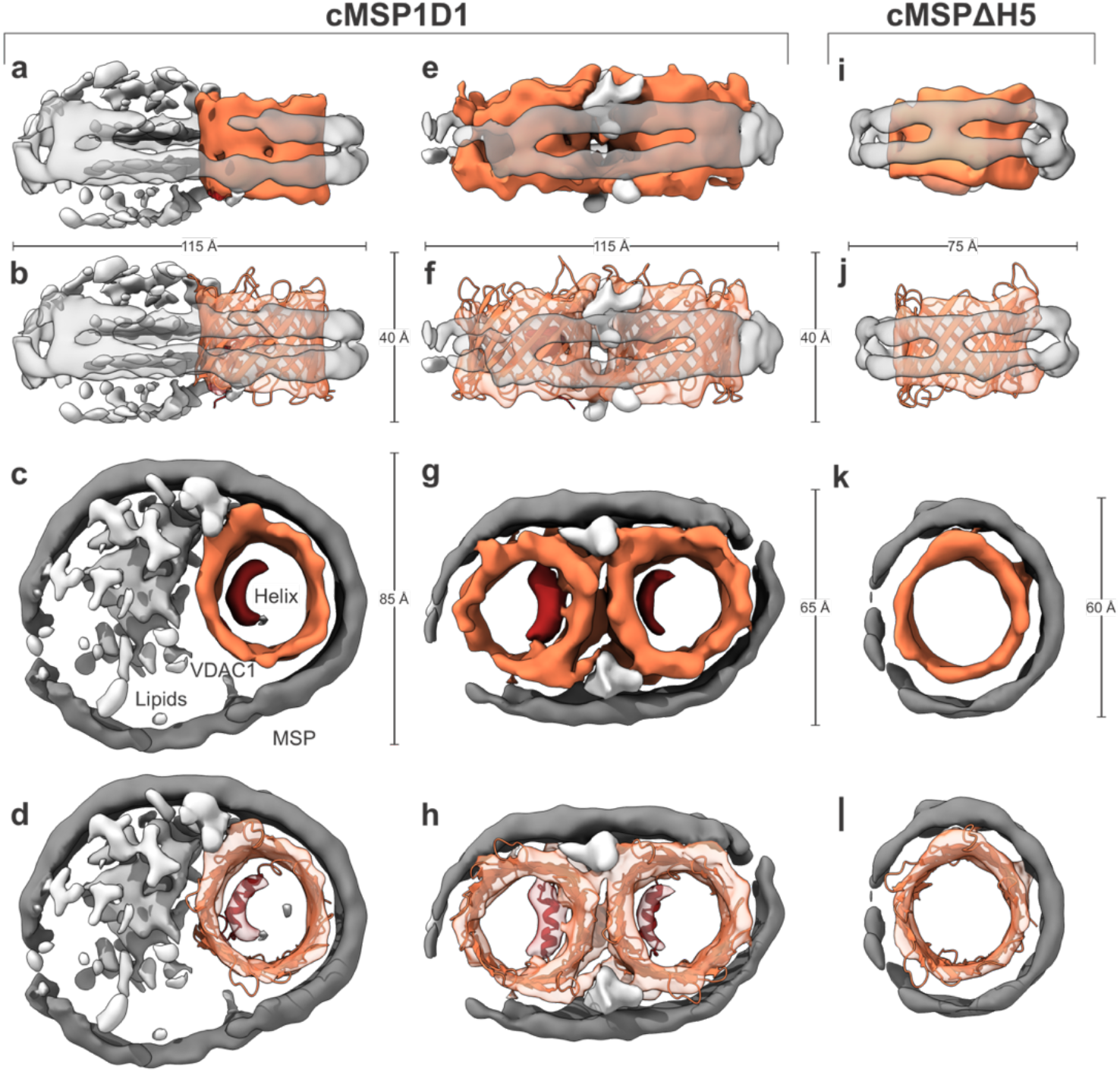
Cryo-EM of VDAC1 reveals different conformational states of its N-terminal helix. Reconstruction of monomeric (**a-d**) and dimeric VDAC1 (**e-h**) in cMSP1D1 lipid nanodiscs of ∼11 nm diameter. (**i-l**) Reconstruction of VDAC1 reconstituted in cMSPΔH5 lipid nanodiscs where VDAC1-N is absent inside the pore (**k**,**l)**. MSP is colored dark gray, lipid noise in light gray, VDAC1 in orange, and the internal N-terminal α-helix in red.

We then set out to characterize VDAC1 in cMSP1βH5 nanodiscs (9 nm diameter), repeating the same strategy as for VDAC1 in 11 nm nanodiscs (**Fig. S4, lower panel**). Gel filtration revealed a single peak roughly at the same volume as the dimers in cMSP1D1 together with a high molecular weight peak, most likely belonging to protein-lipid aggregates (**Fig. S3a**).

Subsequent cryo-EM analysis of the lower molecular weight SEC peak revealed several populations. Roughly 20 % of the particles are in the canonical, helix-inserted monomeric state with a slightly depleted lipid content (**Fig. S3f, Fig. S4**). However, as verified in several independent reconstitutions, most particles (∼65 %) formed smaller nanodiscs with VDAC1 tightly interacting with the MSP of the nanodisc with no lipids visible (**Fig. 2k**). Interactions between the MSP and an incorporated membrane protein have been described previously^55^. While VDAC1 in those “tight” nanodiscs shows an expected inner diameter of ∼3 nm, 15 % of the particles contain VDAC1 with larger pore diameters, ranging up to 6 nm (**Fig. S3f**). Due to their rarity and heterogeneity, particles belonging to those classes were excluded from further processing.

In contrast to the reconstructions of VDAC1 in cMSP1D1 nanodiscs, where the N-terminal helix was clearly resolved within the pore already at the stage of the initial *ab-initio* reconstruction, the VDAC1 reconstruction in cMSP1ΔH5 (9 nm) nanodiscs showed only residual noise at the expected position of the N-helix within the pore (**Fig. 2k**). These observations strongly support that in this particle subset, the structural state of the internal helix is substantially changed as compared to the canonical VDAC1 structure. The absence of N-terminal helix density suggests that it is not stably attached to the β-barrel but rather flexible, most-likely becoming exposed to the channel exterior and thus accessible for interaction partners. This structural data is consistent with the biochemical PM40-modification experiments shown in **Figs. 1c,e**, suggesting helix exposure in the smaller nanodisc. In the exposed state, not only the N-terminal helix but also the β-barrel becomes more flexible, as demonstrated by MD simulations using VDAC1 starting structures with the N-terminal helix inside or outside the β–barrel, respectively (**Fig. S5**). An increased degree of flexibility of the VDAC1 β-barrel was previously shown *in silico* if the N-terminal α-helix was removed^56^.

### BclxL binds to the exposed N-terminal α-helix of VDAC1

After having demonstrated that VDAC1 can expose its N-terminal α-helix, we next wondered whether this structural element might be the interaction site with partner proteins, such as BclxL, as previously suggested using VDAC1 peptides^33^. To address this question, we produced VDAC1 in lipid nanodiscs of different sizes as described above and performed interaction studies with ^2^H,^15^N-labeled BclxL using 2D-[^15^N,^1^H]-TROSY NMR experiments (**Fig. 3**). If the VDAC1 binding epitope is exposed, we expect a change in the NMR spectrum of BclxL. In these experiments, VDAC1 in nanodiscs of 8 nm in diameter induced strong effects on the BclxL NMR spectrum, such as chemical shift perturbations (CSPs) and line broadening (**Fig. 3a**). In contrast, VDAC1 in larger nanodiscs (10 nm) resulted in only very weak effects at the concentrations used in the NMR experiment (identical NMR spectra in **Fig. 3b**). The NMR spectrum of BclxL saturated with a high-affinity BH3 peptide derived from the BH3-only protein PUMA^57^ was not affected by VDAC1 addition, even if 8 nm nanodiscs were used (**Fig. 3c**). These data imply that VDAC1-N binds to the BH3 binding groove of BclxL. This assumption was corroborated by the clustering of the NMR effects shown in **Fig. 3a** at the BH3 binding groove of BclxL (**Fig. 3d**) where BH3 peptides interact (e.g. BAK BH3^58^ shown in blue in **Fig. 3d**).

**Fig. 3.**
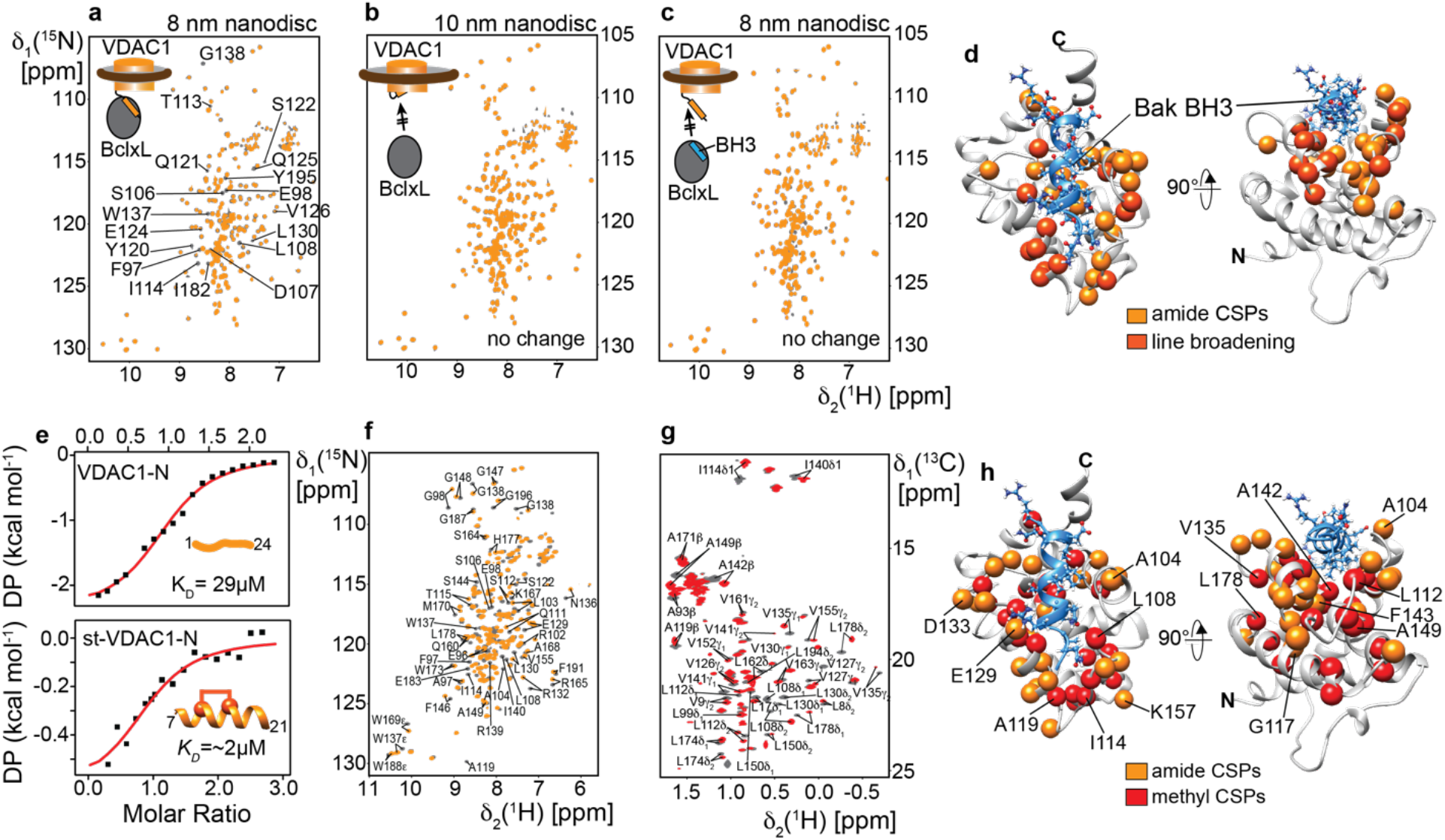
VDAC1 N-terminal α-helix interacts with anti-apoptotic BclxL. (a)-(c) NMR titration experiments with isotope-labeled BclxL (dark grey spectrum) with or without VDAC1 in lipid nanodiscs (orange spectrum). (a) VDAC1 in 8 nm nanodiscs induces chemical shift perturbations (CSPs) in BclxL. A selection of affected signals is labeled in the spectrum. (b) VDAC1 in 10 nm nanodiscs does not show binding to BclxL. (c) BclxL saturated with a high-affinity PUMA BH3 peptide does not interact with VDAC1 in 8 nm nanodiscs, indicating a competitive binding scenario. (d) NMR CSPs and NMR line broadening effects from (a) mapped onto the structure of BclxL. These effects cluster around the canonical BH3 binding site of BclxL. A bound Bak-BH3 peptide is depicted in blue, highlighting that VDAC1-N binds to the canonical BH3 binding site of BclxL. (e) Isothermal titration calorimetry (ITC) with BclxL and a VDAC1 N-terminal peptide (VDAC1-N, top panel) or a hydrocarbon-stapled VDAC1 peptide (st-VDAC1-N, staple between residues 11 and 15, lower panel), indicating a medium to low µM binding affinity. (f) ^2^H,^15^N-labeled BclxL and (g) specifically Ile, Val, Leu, Ala-^1^H,^13^C-methyl labeled BclxL show strong CSP effects upon the addition of st-VDAC1-N. (h) The CSP effects in (f) and (g) cluster to the BH3 binding site of BclxL, suggesting that VDAC1-N is the main interaction site with BclxL.

To monitor the structural state of VDAC1 in 8 or 10 nm nanodiscs by NMR we recorded 2D-TROSY spectra with ^2^H,^15^N-labeled VDAC1 (**Fig. S6**). While the NMR signals are well resolved in the larger nanodisc (**Fig. S6, left**), strong line broadening can be seen in the smaller nanodisc (**Fig. S6, right**), indicative of pronounced intrinsic motions in the µs to ms time scale, presumably induced by the tight packing of the nanodisc scaffold protein around VDAC1, as seen in the cryo-EM data (**Fig. 2**).

Since the main difference between the investigated VDAC1 samples is the degree of N-terminal α-helix exposure, we next wanted to confirm whether this structural element in VDAC1 is indeed the main interaction site with BclxL. For this, we first used a VDAC1 N-terminal peptide (residues 1-24), representing the entire N-terminal part which adopts a random coil structure in solution (**Fig. S7**). The binding affinity between this peptide and BclxL was evaluated with isothermal titration calorimetry (ITC) yielding a *K*_*D*_ value of 29 µM (**Fig. 3e**, top panel). All BH3 peptides that interact with the binding groove in BclxL adopt an α-helical secondary structure. The VDAC1 N-terminus is also α-helical when bound to the pore interior. To mimic the bound structure, we used an optimized hydrocarbon-stapled VDAC1 peptide encompassing residues 7-21 (st-VDAC1-N). This peptide adopts an α-helical secondary structure, as confirmed by CD spectroscopy (**Fig. S7**). It binds more tightly to BclxL (*K*_*D*_ of ∼2 µM) than the linear peptide (**Fig. 3e**, lower panel), which could also be confirmed by NMR titrations (**Fig. S8**). This concept has previously been applied to BH3 peptides to obtain more efficient activators of pro-apoptotic Bcl2 proteins ^59^. We used the higher affinity st-VDAC1-N for NMR titrations monitored by the backbone amide (**Fig. 3f**) or sidechain methyl group signals (**Fig. 3g**) in BclxL as probes for binding. Both groups of resonances in BclxL undergo strong CSPs upon peptide binding. Mapping of the CSPs on the structure of BclxL clearly shows that the st-VDAC1-N peptide specifically interacts with the BH3 binding groove of BclxL (**Fig. 3h**), consistent with the NMR results with full-length VDAC1 in lipid nanodiscs (**Fig. 3d**). 2D NMR titration experiments with ^15^N-labeled VDAC1-N and unlabeled BclxL confirm the interaction (**Fig. S9a**) and show that the central α-helical part of VDAC-N that has a slight homology to BH3 peptides (**Fig. S9b**) interacts with BclxL (**Fig. S9c**).

### High-resolution structure of BclxL in complex with the VDAC1 N-terminal α-helix

To obtain a higher resolution picture of the VDAC1-N - BclxL interaction, we next screened for suitable expression constructs for X-ray crystallography. To ensure that the VDAC1-N peptide remains stably bound to BclxL we produced a single chain construct where VDAC1-N (1-26) is fused to the C-terminus of BclxLΔLT (lacking its flexible loop and the transmembrane helix after residue 209) (**Fig. 4a**). This construct allowed for crystal formation of suitable quality for a high-resolution structure determination by X-ray diffraction. These data led to the structure determination of BclxLΔLT in complex with the VDAC1-N peptide at 1.95 Å resolution (**Fig. 4b, Table S1**). The parallel relative orientation of BclxL and the VDAC1-N peptide in the complex structure results in the C-termini of both proteins being located at proximal positions next to the membrane surface (**Fig. 4c**).

**Fig. 4.**
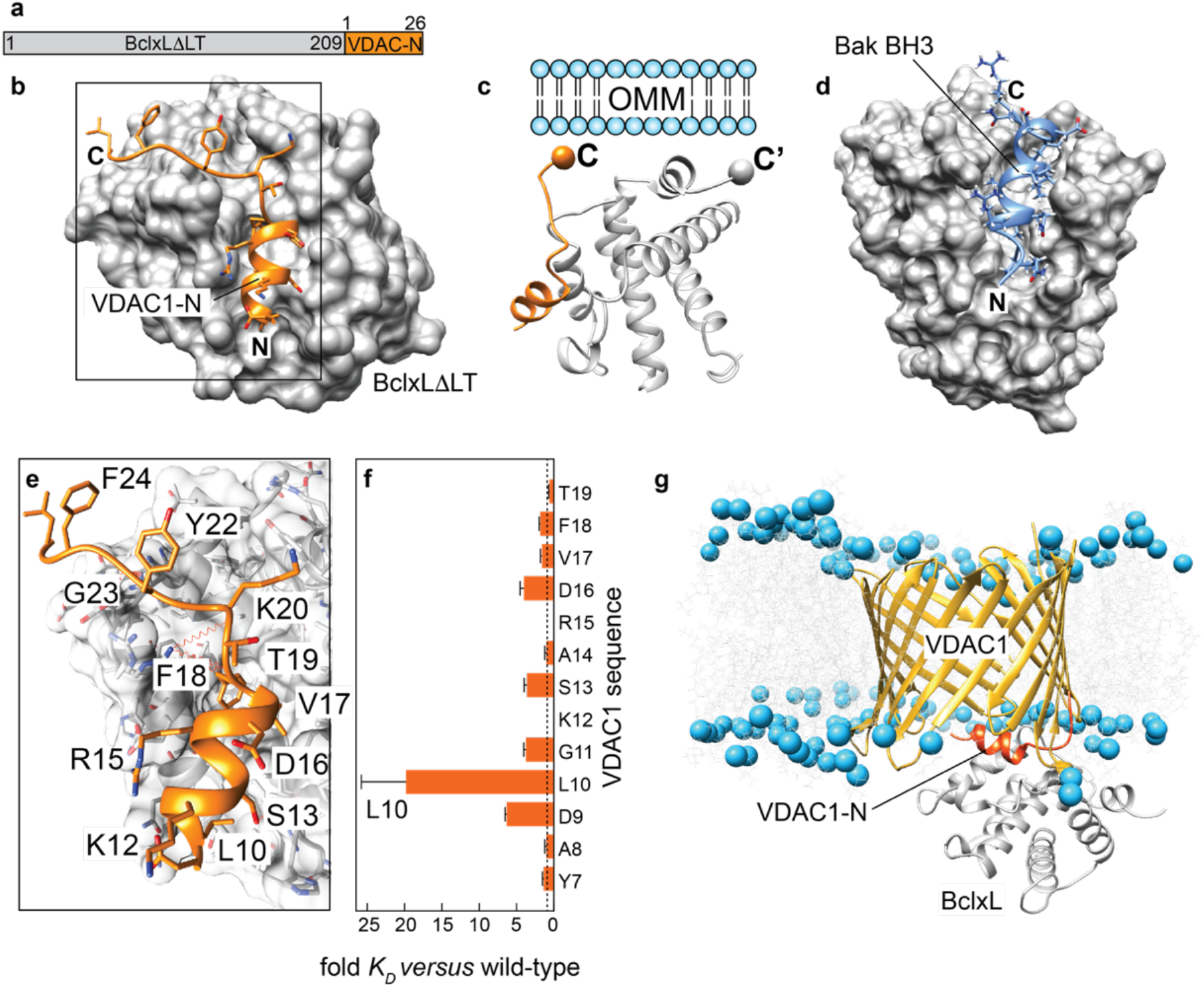
Complex structure of BclxL and VDAC1-N. (a) Single-chain construct to enable structure determination of the VDAC1-N-BclxL complex. (b) Structure of BclxLΔLT in complex with VDAC1-N determined here. (c) Schematic view of the C-termini of VDAC1-N and BclxL, both oriented in the same direction. (d) Complex structure of BclxL and the BH3 domain of Bak^58^. (e) Close-up view of the interface between BclxL and VDAC1-N in the complex. Amino acids in VDAC1-N participating in the interaction are labeled. (f) Relative affinity of VDAC1 peptides in an alanine scanning experiment. Substitution of Leu10, facing the BclxL binding groove shows the most pronounced effect. (g) Complex structural model of the VDAC1 and BclxL based on the structural data reported in this study. VDAC1-N is shown in orange, lipid phosphate groups are visualized as blue spheres.

In the full-length protein, the C-terminal end of VDAC1-N is attached to its β-barrel and the C-terminus of BclxLΔTM to its transmembrane helix, respectively. To enable the formation of the herein determined complex structure, VDAC1 and BclxL must have a suitable orientation in the membrane. The soluble domain of BclxL is known to be located at the cytosolic side of the OMM, like all other Bcl2 proteins^60^. For VDAC1, the orientation in the OMM has been probed by a caspase reporter assay in intact cells^61^. It showed that both N- and C-termini are facing the mitochondrial intermembrane space, suggesting that the attachment point of VDAC1-N at the β-barrel is on the cytosolic side. Once VDAC1-N becomes detached from the pore interior, it can swing out toward the cytosol – an orientation that is consistent with MD simulations with our structural model (**Fig. S5a**). Even though the general binding mode between BclxL and VDAC1-N is similar to BH3 peptides, as shown in the BclxL-Bak-BH3 complex^58^, the exact binding site of VDAC1-N is shifted toward the periphery of the BclxL binding groove. Furthermore, the C-terminal end of VDAC1-N binds to BclxL in an extended conformation, which was not seen for BH3 peptides so far. VDAC1-N forms an amphipathic helix with the mostly hydrophobic side chains of Leu10, Ser13, Asp16, Val17 and Phe18 participating in binding (**Fig. 4e**).

To validate the structural model of the complex, we performed alanine scanning experiments where every individual residue in the binding region of VDAC1-N was mutated to alanine (**Table S2**), followed by NMR-detected affinity measurements with each peptide (**Fig. 4f, Fig. S10**). These experiments showed a strong effect on the binding affinity for residues that are oriented toward BclxL in the complex, with Leu10 having the biggest impact. Leu10 is the only large hydrophobic residue in VDAC1-N that is establishing a hydrophobic interaction with the BH3 binding groove in BclxL, thus markedly contributing to the overall binding affinity. Leu10 is also one of the main interaction sites between VDAC1-N and the β-barrel. Consequently, its mutation to cysteine weakens the helix attachment and enhances its exposure (**Fig. S1b-e**).

We next set out to assemble a structural model of the full-length VDAC1-BclxL complex *in silico* by attaching the experimental BclxL-VDAC1-N structure to the VDAC1 β-barrel. Initial structural clashes in the assembled structure were resolved in a 200 ns molecular dynamics simulation. In the resulting structural model (**Fig. 4g**), the dissociated VDAC1-N is located slightly outside the β-barrel where it can interact with the BH3 binding groove of BclxL. A recent NMR study of full-length BclxL in lipid nanodiscs showed that the BH3 binding groove of the soluble domain of BclxL is facing toward the membrane surface^62^, which resembles the orientation of BclxL in the complex with VDAC1.

### The VDAC1 N-terminus acts as a sensitizer BH3 protein to inhibit anti-apoptotic BclxL

Next, we aimed at addressing the functional relevance of the VDAC1-BclxL complex in the induction of apoptosis via mitochondrial outer membrane permeabilization (MOMP). In recent literature reports, it has been postulated that VDAC1 oligomers alone can form large pores to enable MOMP^21^ and even allow for the exit of mitochondrial DNA^22^. To address this question, we conducted *in vitro* experiments with VDAC1 in liposomes loaded with cytochrome C or lysozyme that are both of similar size (12 and 14 kDa, respectively) (**Fig. S11a, b**). Such VDAC1 proteoliposomes and liposomes without VDAC1 were subjected to a size exclusion chromatography column and the remaining proteins inside the liposomes were quantified by their corresponding SDS-PAGE band intensity. For both proteins, no translocation across VDAC1 could be detected. However, the flux of ATP across VDAC1 in proteoliposomes was possible in the same setup (**Fig. S11c**), indicating that VDAC1 was functional but did not form a larger pore in liposomes that would allow for the translocation of proteins.

Thus, we wondered whether VDAC1 could rather have an indirect pore-forming activity via its interaction with Bcl2 proteins. As a well-established model system, we used the pore-forming Bcl2 protein Bak that can be efficiently inhibited by anti-apoptotic BclxL^63-65^. To probe the pore forming activity of Bak alone, in presence of BclxL and with increasing amounts of the VDAC1-N peptide, we utilized an established liposome pore forming assay^66^. At a rather high concentration of 1 µM, Bak forms pores without activation by BH3-only proteins (autoactivation conditions), which can be partially inhibited by BclxL (**Fig. 5a**, black curve). The addition of the VDAC1-N peptide leads to complex dissociation via competitive binding to BclxL and consequently to a concentration dependent increase in the pore forming activity of Bak (**Fig. 5a**, brown to yellow curve, **Fig. 5b**) to the level of Bak without BclxL present (red curve).

**Fig. 5.**
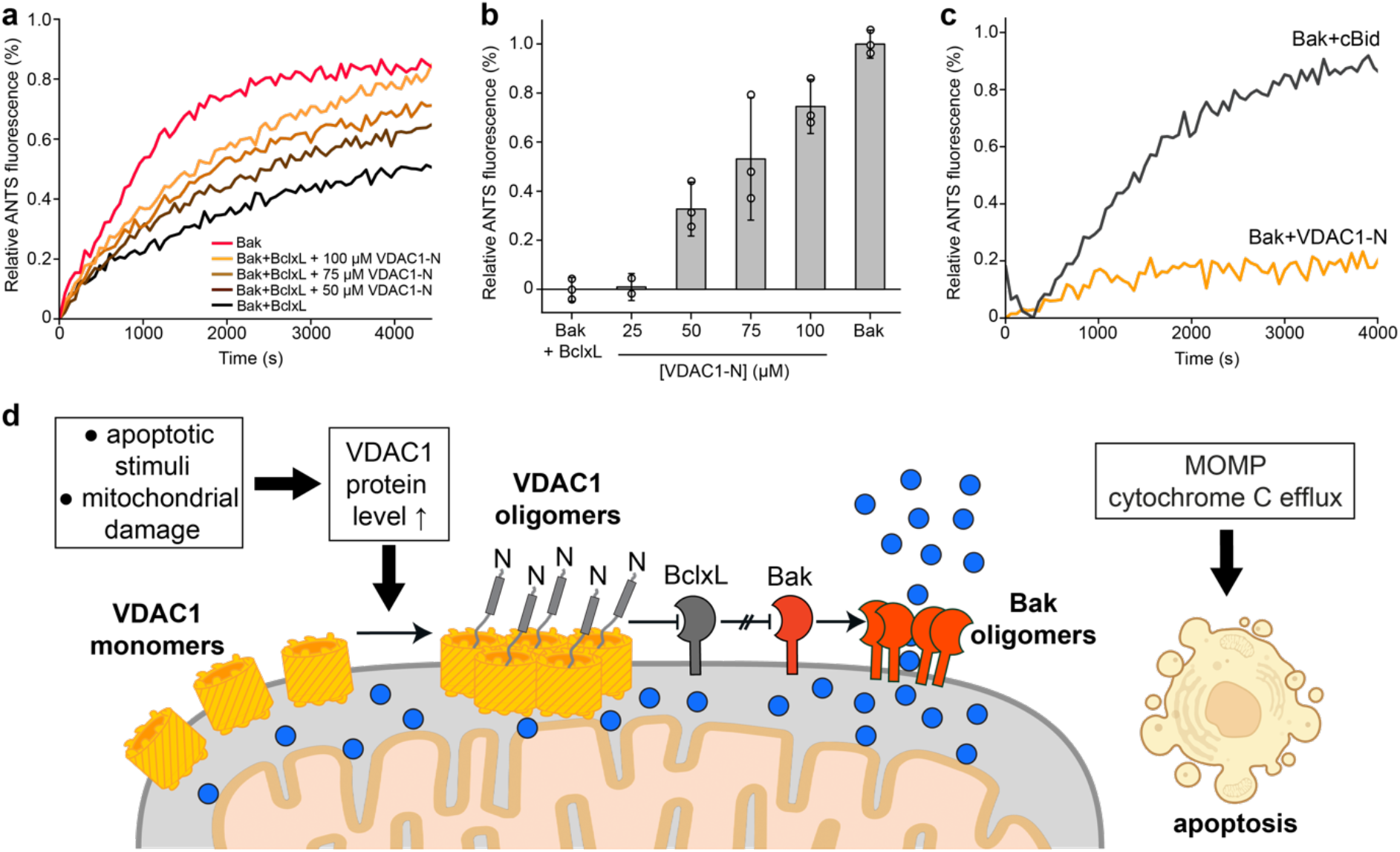
VDAC-N acts as a sensitizer BH3 protein to induce Bak mediated membrane permeabilization. (a) Pore formation of BakΔTM (1 µM concentration, red) is inhibited by the addition of BclxLΔTM (500 nM, black). The addition of VDAC1-N (25-100 µM) induces Bak pore formation in the presence of BclxL in a concentration-dependent manner (brown to yellow). (b) Fluorescence intensity taken from panel (a) averaged between 4000 and 4500 s. The standard deviation (error bars) was calculated from at least two technical replicates. (c) Pore formation of 200 nM BakΔTM is activated by the addition of 50 nM cBid (black) but not by VDAC1-N (yellow, 40 µM concentration). (d) Functional model: apoptotic stimuli or mitochondrial damage lead to VDAC1 oligomerization, inducing the exposure of its N-terminal α-helix which can neutralize anti-apoptotic Bcl2-like proteins, such as BclxL, to induce the execution of MOMP by Bak. Blue spheres: cytochrome C.

To exclude a direct off-target effect of VDAC1-N on the activity of Bak, we tested whether VDAC1-N can activate Bak pore formation in comparison with the known Bak activator cBid^65^. The results in **Fig. 5c** clearly demonstrate that even a large excess of VDAC1-N (200 nM *versus* 40 µM) does not lead to Bak activation whereas cBid can already fully activate Bak at a sub-stoichiometric concentration of 50 nM. These results show that VDAC1-N promotes Bak pore formation by neutralizing the inhibitor BclxL. This behavior is reminiscent of sensitizer BH3 proteins, such as BAD or NOXA, that exhibit pro-apoptotic activity by neutralizing the anti-apoptotic members^3^.

## Discussion

In this study we show that VDAC1 can expose its N-terminal α-helix to the cytosolic side of the mitochondrial outer membrane, which is a prerequisite to interact with the anti-apoptotic Bcl2 protein BclxL. We present a thorough set of biophysical and structural data that confirm a specific interaction between VDAC1-N and BclxL. Via its interaction with BclxL, VDAC1 can act as a so-called sensitizer BH3 protein, a class of pro-apoptotic Bcl2 proteins that inhibit the anti-apoptotic Bcl2 family members and eventually induce the mitochondrial apoptosis pathway^3^ (**Fig. 5d**). In our liposome pore forming assays (**Fig. 5a-c**), we clearly see that the BclxL-mediated inhibition of the pore-forming activity of the Bcl2 protein Bak is released in a dose-dependent manner upon the addition of VDAC1-N. Thus, the exposure of the VDAC1 N-terminal helix can be considered a key event that is required to transduce its pro-apoptotic functionality. In full agreement with our model, VDAC1 where the N-terminal helix was deleted (Δ26-VDAC1) was not able to induce apoptosis and no cytoprotective effect of anti-apoptotic partner proteins, such as BclxL, could be observed^34^.

Under non-apoptotic conditions, the N-terminal helix is stably attached to the pore interior, as demonstrated by the available structures of VDAC1^8-10^. Free energy calculations with the canonical VDAC1 structure indicate that the helix-inserted state is strongly favored^51^. Therefore, a trigger is necessary to induce such a substantial structural change within VDAC1. Our data suggest that VDAC1 oligomerization is the key event that induces the release of the N-terminal helix (**Fig. 1**). VDAC1 oligomers^20,23^ have been reported to be induced by diverse stimuli, such as VDAC1 up-regulation^67^, changes in the mitochondrial lipid composition^24,25^, increased Ca^2+^ levels^26,27^, low pH^28^, oxidative stress^22,29^ or specific small molecules that inhibit the respiratory chain^68^. VDAC is highly abundant in the mitochondrial outer membrane (MOM)^69^ and has been shown to form dynamic supramolecular assemblies of various sizes ranging from monomers up to 20-mers^20^. Thus, slight changes in the MOM environment and in the VDAC protein levels have the potential to strongly modulate the VDAC oligomeric state. Even though the mechanistic details leading to tight VDAC oligomerization remain unclear, there is culminating evidence that VDAC oligomerization is a marker for apoptosis in human cells^2,21^. This interesting feature suggests that VDAC might be an internal checkpoint of the functional state of mitochondria, which contrasts with the action of pro-apoptotic Bcl2 proteins where apoptosis signals originate from outside the cell, the cytoplasm or other organelles^70^. Moreover, because of the role of VDAC in apoptosis induction it is considered an emerging drug target in cancer therapy^69,71^.

We observed strong VDAC1 oligomerization in the negatively charged detergent cholate but not in the structurally very similar zwitterionic detergent CHAPS or the structurally distinct zwitterionic detergent LDAO (**Fig. 1b**). In these experiments the degree of oligomerization highly correlated with the exposure of the N-terminal helix as probed by chemical modification experiments with PM40 (**Fig. 1c**). In line with these observations, enhanced helix exposure was also detected in liposomes assembled with negatively charged lipids (**Fig. S1d**,**e**). This finding is consistent with previous data showing an interaction between VDAC1-N and a negatively charged lipid^49-51^ or detergent surface^72^. Of particular importance, VDAC1 in lipid nanodiscs of 8 and 10 nm diameters^46,47^ showed strong differences in the exposure of the N-terminal helix, even though a VDAC1 monomer was mainly present in each nanodisc (**Fig. 1d,e**). In the smaller (8 nm) nanodiscs, the exposure of VDAC1-N was more pronounced than in the larger (10 nm) nanodisc. Structural insights obtained by cryo-EM demonstrate that only 20% of the VDAC1-cMSP1βH5 particles adopt the canonical, helix-inserted state, whereas this is the case for all particles with VDAC1 in the larger nanodiscs (**Fig. 2, Fig. S3**).

Since the “tight” VDAC1 state in cMSP1ΔH5 nanodiscs is stabilized by protein-protein interactions with the MSP instead of a lipid environment, we consider it very likely that this setup mimics the oligomeric state where protein-protein interactions between VDAC1 protomers dominate^20^. This assessment is further supported by the very similar outcome of the PM40 modification experiments in detergent micelles, liposomes and small lipid nanodiscs (**Fig. 1, Fig. S1**). Furthermore, our nanodisc setup shows that VDAC1 can adopt a helix-exposed structural state with the β-barrel fully intact (**Fig. 2**) and which is capable of binding to the partner protein BclxL (**Fig. 3**). Thus, we consider the chosen nanodisc strategy functionally relevant and necessary to obtain further structural information. Moreover, nanodiscs have been previously used to selectively stabilize functional structural states of membrane proteins^73,74^. By using circularized MSP^41,75^ we obtained stable VDAC1 preparations in 9 and 11 nm nanodiscs with optimized homogeneity (**Fig. S3**). This setup facilitated the structural characterization of both states by single particle cryo-EM. Despite recent progress in cryo-EM, membrane proteins such as VDAC1 with a molecular weight of just 31 kDa without a large protein mass outside the membrane are still very challenging^76^.

Nonetheless, we achieved a resolution range of 6.5 to 7.8 Å sufficient to detect the location of VDAC1-N in the helix-inserted canonical state in 11 nm nanodiscs and its absence in the exposed state in the smaller 9 nm nanodiscs. Even though a cryo-EM resolution of ∼6 Å leaves some degree of uncertainty, it is most likely that VDAC1-N in 9 nm nanodiscs is either extruded from the pore or substantially more mobile than in the canonical structures. In addition, the strong variation of noise surrounding the particle suggests that the exposed N-terminal helix is tumbling outside the pore in a dynamic manner, in line with our MD simulations (**Fig. S5**). Furthermore, we identified a dimeric VDAC1 state in 11 nm nanodiscs (**Fig. 2**) with the dimerization interface located at that side of the β-barrel wall where the N-terminal helix is attached, resembling a previously published crystal structure^54^. The decent resolution in our cryo-EM structure was likely enabled by the specific interaction between VDAC1 and the MSP forming the rim of the nanodisc particles. Interactions of the MSP with membrane proteins in cryo-EM structures have been previously analyzed^55^ and the interaction with the MSP also enabled the design of water-soluble MSP-membrane protein fusions in *E. coli*^77^.

Although controversially discussed in the literature^17^, previous studies have suggested the direct involvement of VDAC1 in forming a large pore in the outer mitochondrial membrane^16,22,30^. Our data does not support such a direct role of VDAC1. Nonetheless, in our single particle cryo-EM analysis we observed a minor species (15%) with a larger pore diameter of up to 6 nm (**Fig. S3f**), which was excluded from further analysis due to heterogeneity. Even though the relevance and nature of a larger VDAC1 species is unclear, we can conclude that such larger VDAC1 pores can be formed under conditions where the β-barrel is destabilized.

Our data presented here support a model where, triggered by oligomerization, VDAC1-N dissociates from the interior surface of the β-barrel. The availability of VDAC1-N at the MOM surface enables its interaction with anti-apoptotic BclxL, which becomes inhibited by binding of VDAC1-N to its canonical BH3 groove. Due to the high abundance of VDAC1 in the MOM^69^, the µM affinity between VDAC1-N and BclxL would be insufficient to prevent this interaction. Looking at apoptosis induction mechanisms at the mitochondrial surface, the action of VDAC1-N is reminiscent of sensitizer BH3 proteins with pro-apoptotic activity^3^. Thus, VDAC1 can be regarded as an indirect inducer of cytochrome C release and apoptosis by acting as a sensitizer BH3 protein (**Fig. 5d**). In agreement with the presented functional model, an interplay between VDAC and the pro-apoptotic Bcl2 proteins Bak and Bax in forming membrane pores in liposomes and mitochondria has been reported previously^30^ and deletion of VDAC-N was reported to abolish its apoptotic functionality^34^.

In a more general context, our study suggests a novel regulation mechanism involving β-barrel membrane proteins where the plasticity of the β-barrel enables the occupation of different functional states. Such a mode-of-action is sensitive to factors like oligomerization or partner protein and metabolite binding and offers an additional layer of regulation, as shown recently for a chloroplast metabolite channel^78^.

## Acknowledgements

This study was supported by the Helmholtz Society (to F.H. VG-NG-1035) and the German research foundation (DFG) within the CRC1035 (project B13, project number 201302640 to FH). We acknowledge spectrometer time at the Bavarian NMR Center (www.bnmrz.org), Drs. Elisabeth Häusler and Shakeel Shahid for initial experiments, and Drs. Gerd Gemmecker and Sam Asami for general NMR support, Marie Tran for help with protein production and Dr. Oleg V. Maltsev for peptide synthesis. We acknowledge the use of the X-ray Crystallography Platform at Helmholtz Munich. The cryo-EM data were collected at “Cryo-EM SoN”, the cryo-EM infrastructure of the University of Münster, funded by the DFG – Project number 496113311. C.G. acknowledges funding through CRC1430, Project A04 (DFG, 424228829) and CRC1348, Project A15 (DFG, 386797833).

## Author contributions

M.D, U.G., G.B, R.J., K.L., K.F. and F.H. conducted research and analyzed data. M.D., U.G., G.B., C.G., R.J. and F.H wrote the manuscript. All authors commented on and approved the manuscript. F.H. designed the study. D.N., C.G. and F.H acquired funding.

## Data availability

The structural coordinates of the X-ray structure of BclxLΔLT bound to the VDAC1-N peptide were deposited at the RCSB databank (accession code 9HPS). The cryo-EM maps of monomeric VDAC1 in cMSPD1, dimeric VDAC1 in cMSPD1 and monomeric VDAC1 in cMSPΔH5 have been deposited to the Electron Microscopy Data Bank (EMDB) under the accession codes EMD-XXX, EMD-XXX, and EMD-XXX, respectively. The respective cryo-EM datasets have been deposited to EMPIAR under accession codes EMPIAR-XXX, EMPIAR-XXX, EMPIAR-XXX.

