## Supplemental Information for "Structural basis of apoptosis induction by the mitochondrial voltage dependent anion channel"

### Supporting Information Methods

#### Molecular cloning and protein production

The DNA for human BclxL where the transmembrane helix (TMH) alone or the TMH and the flexible loop (residues 44-84) was deleted, BclxLΔTM or BclxLΔLT, respectively, were cloned into a pET21a plasmid (Merck). VDAC1, inserted into pET21a, was produced, refolded and purified as described[1]. Site directed mutagenesis of BclxL or VDAC1 was done with the Quikchange Lightning Kit (Agilent).

The voltage-dependent anion channel 1 (VDAC1) and variants were produced in *E. coli* BL21(DE3) by induction with 1 mM IPTG. Expression was carried out for 3-4 hours at 37 °C. After cell lysis and centrifugation for 20 min at 40 000 g, the pellet was resuspended in lysis buffer (50 mM Tris pH 7.0, 250 mM NaCl, 20 mM BME) + 1% TritonX-100. The mixture was then subjected to centrifugation at 40 000 g. Subsequently, resuspension and centrifugation were repeated for two more times with lysis buffer without TritonX-100. The pellet was then solubilized in IMAC buffer (6 M GdmCl, 50 mM Tris pH 8.0, 100 mM NaCl) overnight. After another round of centrifugation, the supernatant was applied to a Ni-NTA gravity follow column (Cytiva). The column was washed with IMAC buffer and the protein then eluted using IMAC buffer + 500 mM imidazole. The protein was then dialyzed against 20 mM Tris pH 7.5, 50 mM NaCl, 1 mM EDTA and 10 mM BME. The precipitated protein was solubilized in 6 M GdmCl, 100 mM NaPi pH 7.0, 100 mM NaCl, 1 mM EDTA and 5 mM DTT and the concentration adjusted to 5 mg/mL. VDAC1 wild-type and variants were refolded at 4 °C by dropwise addition of the protein into a 10x volumetric excess of refolding buffer (25 mM NaPi pH 7.0, 100 mM NaCl, 1 mM EDTA, 5 mM DTT and 1% LDAO (Avanti Polar Lipids)). The mixture was then stirred for 6-8 hours and then dialyzed overnight against 10 mM NaPi pH 6.5, 1 mM EDTA, 100 mM NaCl and 3 mM DTT. Subsequently, size-exclusion chromatography was conducted by applying the protein to a HiLoad™ 16/600 Superdex™ 200 pg column (Cytiva). The column was equilibrated in 10 mM NaPi pH 6.5, 100 mM NaCl, 1 mM EDTA, 5 mM DTT and 0.1% LDAO. Monomeric protein was pooled and concentrated to ~300 μM.

To produce $^{2}H,^{15}N-CH_3\varepsilon-^{13}C$-methionine-VDAC1-V3M, protein expression was conducted in M9 medium containing 99% $D_2O$, 1 g/L $^{15}N-NH_4Cl$ and 2 g/L deuterated glucose. 70 mg/L L-methionine (ε-methyl-$^{13}C$) (Sigma-Aldrich) was added to the M9 expression medium 1 h before induction of protein expression with 1 mM IPTG. After 4 hours of shaking at 37°C, cells were harvested by centrifugation. $^{2}H,^{15}N-CH_3\varepsilon-^{13}C$-methionine-VDAC1-V3M was purified as described for wild-type VDAC1[1-3].

Plasmids encoding for cBid and the soluble domains of Bak (BakΔTM, residues 1-186) and BclxL (BclxLΔTM, residues 1-212, BclxLΔLT, residues 1-43, 85-212 and BclxLΔLT fused to VDAC1-N: BclxLΔLT-LEGG-VDAC1-N(1-26)-GG-$His_6$) were expressed in *E. coli* as previously described [4-7]. In brief, *E. coli* BL21 (DE3) cells were transformed with the respective plasmids and grown at 37°C. For BakΔTM, protein expression was induced with 0.1 mM IPTG and grown for 16-20 h at 20°C, whereas for cBid and BclxL, protein expression was induced with 1 mM IPTG and grown for 4 h at 37°C. In the resulting cBid protein, a thrombin cleavage site is replacing the original caspase-8 site [6]. Purification of cBid, BakΔTM, and BclxL was performed as previously described [6]. Briefly, cell pellets were resuspended in lysis buffer (50 mM Tris pH 8, 200 mM NaCl, 1 mM EDTA) supplemented with 2 mM PMSF and 1 mg/mL lysozyme. The suspension was then subjected to sonication for 30 min (1-s pulse, 3-s pause, 30% amplitude), incubated together with DNase1 and 5 mM $MgCl_2$, and the cell debris was removed via centrifugation. The supernatant was applied to a Ni-NTA gravity flow column equilibrated with buffer A (20 mM Tris pH 8.0, 100 mM NaCl, 5 mM BME), washed with 20 CV buffer A, 20 CV buffer A supplemented with 10 mM imidazole followed by elution with 20 CV buffer A supplemented with 400 mM imidazole. The elution fraction was dialyzed into buffer A overnight and for cBid, 20 U thrombin/L cell pellet was added. 5 % glycerol was added to protein solutions before concentrating for size exclusion chromatography (SEC) on an ÄKTA Pure system equipped with a HiLoad 16/600 Superdex 75 pg column equilibrated with 20 mM NaPi pH 7.0, 50 mM NaCl, 0.1 mM EDTA and 5 mM BME.

Constructs encoding GB1-VDAC peptides were cloned into an in-house modified pET vector encoding GB1 with an N-terminal $His_6$ tag and TEV site followed by a thrombin site ($His_6$-TEV-GB1-thrombin-peptide). We produced the following constructs: GB1-VDAC1-N (residues 2-25) and GB1-VDAC2-N (residues 2-36). Expression and purification of the GB1-VDAC peptides was described previously[8]. Following SEC, the GB1-VDAC peptides were digested with TEV (1:40 molar ratio) in the presence of 5 mM BME overnight at 4°C. Reverse

Ni-NTA was performed to isolate the cleaved peptides, which eluted mainly during sample loading. Cleavage of the GB1-VDAC peptides was confirmed by SDS-PAGE analysis.

Synthetic VDAC1 peptides were purchased from Bio-Synthesis and Biomatik. Isotope labeled ($^{13}C$ and $^{15}N$) VDAC1 peptides (residues 1-26) were produced recombinantly in *E. coli* as a fusion construct with GB1 ($His_6$-GB1-TEV-VDAC1-N) [8]. Cells were grown in minimal media supplemented with 1g/L $^{15}NH_4Cl$ and 2g/L $^{13}C_6$-glucose until an $OD_{600}$ of 0.7 at 37°C. Protein production was induced by the addition of 1 mM IPTG and cells were shaken for further 4 h at 37°C, followed by harvesting by centrifugation. Protein purification was done by Ni-NTA chromatography, followed by digestion with TEV protease and reverse Ni-NTA chromatography to collect the VDAC1-N peptide in the flow-through and wash fractions, which were dialyzed against $H_2O$, lyophilized and dissolved in NMR buffer (20 mM NaPi, pH7.0, 50 mM NaCl, 0.5 mM EDTA, 5 mM DTT) for further analysis.

**Lipid nanodisc assembly**

Assembly of VDAC1-containg lipid nanodiscs was done according to established protocols[9-12]. VDAC1 and VDAC1-V3M nanodiscs with MSPΔH5, MSP1D1, cMSPΔH5 and cMSP1D1 were assembled with a VDAC1:MSP:lipid ratio of 1:6:40, 1:6:60, 1:6:55 and 1:6:80, respectively. The lipid mixture consisted of 75% DMPC and 25% DMPG. Respective amounts of MSP buffer (20 mM Tris pH 7.5, 100 mM NaCl, 0.5 mM EDTA), lipids in 100 mM sodium cholate, MSP and VDAC1 or VDAC1-V3M in LDAO were mixed and incubated for 1 h at room temperature (RT). Subsequently, the mixture was incubated for 1.5 h at RT with 0.5 g washed Biobeads-SM2 (Bio-Rad) per mL of reaction, while rocking. The beads were then removed by filtration and washed with buffer. The assembled nanodiscs were then subjected to Ni-NTA and a Superdex 16/600 200 pg column (Cytiva). For the preparation of nanodiscs for cryo-EM, the size exclusion chromatography buffer was 20 mM HEPES pH 7.0, 100 mM NaCl, 0.5 mM EDTA and 2 mM DTT.

**Chemical crosslinking and modification of VDAC1**

Protein crosslinking in detergent micelles was performed in 20 mM HEPES-NaOH, pH 7.5, 50 mM NaCl with a final VDAC1 concentration of 20 µM in the presence of 20 mM of different detergents. After a 30 min preincubation at RT, 50x molar excess of the amino-selective crosslinker bissulfosuccinimidyl suberate ($BS^3$, ThermoFisher Scientific, Cat.No. 21580) was applied for 30 min, as reported previously[13]. The reaction was stopped by the addition of 50 mM (final concentration) Tris/HCl pH 7.5 and further incubation for 15 min. For polyethyleneglycol (40 kDa) maleimide (PM40) modification, 20 μM PM40 (Sigma Aldrich) was applied for 15 minutes, and the reaction was stopped by the addition of a large excess of BME. Finally, the samples were analyzed by SDS-PAGE and Coomassie or silver staining. Nanodisc samples were prepared as described above. 20 μM PM40 was applied directly after detergent removal to 20 μM VDAC nanodiscs as indicated above. For the preparation of liposomes, 10 μM VDAC1 mixed with 5 mg/mL lipids (100 % POPC or 50 % POPC + 50 % POPG) solubilized with 0.5 % LDAO and 0.1 % Triton X-100. After 30 minutes preincubation, detergents were removed using two rounds of Bio-Beads-SM2 (200 mg/mL at each round). 50x $BS^3$ or 2x PM40 was applied on liposome samples, which were then 2x flash frozen and thawed on a thermoblock at room temperature. Following 30 min $BS^3$ and 1 h PM40 reaction, the samples were quenched as above and finally analyzed by SDS-PAGE.

**Liposome translocation assay**

10 μM VDAC1 was mixed with 10 mg/mL Egg PC lipids solubilized with 0.5 % LDAO and 0.1 % Triton X-100 in 20 mM HEPES-NaOH, pH 7.5, 100 mM NaCl. After a 30 min preincubation, detergents were removed using 3 rounds of Bio-Beads (2x 100 mg/mL and 1x 200 mg/ml). Molecules (2 mM ATP, 1.5 mg/mL cytochrome C or lysozyme, 1 mg/mL BSA) to be transported were added into liposomes and three rounds of freeze-thaw cycles were applied. Then, the liposomes were passed through 0.2 μm filter for 15 times in a mini extruder (Avanti Polar Lipids) and were injected into a Superose 6 Increase 10/300 GL (Cytiva). Finally, the peak fractions were analyzed using SDS-PAGE and silver staining. ATP was detected by chemiluminescence with an ATP determination kit (Promega, cat# A22066) following the instructions of the manufacturer.

**Liposome permeabilization assay**

The liposome permeabilization assays were performed at 30°C as previously described [14,15]. Briefly, 200 nM BakΔTM, 50 nM cBid and 1-40 μM VDAC-N peptides were used for measurements at low BakΔTM concentrations whereas 1 μM BakΔTM and 500 nM BclxLΔTM were used for measurements under auto-active BakΔTM conditions. VDAC1-N peptide was titrated into the Bak-BclxL complex from 20 to 100 μM under auto-active BakΔTM conditions. The liposomes were prepared with the following lipid composition (mass percentage) to mimic the outer mitochondrial membrane (OMM), as previously described{Kale, 2014 #727}. Briefly, the OMM-like lipids were prepared by combining 38% L-α-phosphatidylcholine (PC), 25% L-α-phosphatidylethanolamine (PE), 10% L-αphosphatidylinositol (PI), 10% 1,2-dioleyl-sn-glycero-3-phospho-L-serine (DOPS), 7% 1,1′,2,2′-tetra-(9Z-octadecenoyl)cardiolipin and 10% 18:1 DGS-NTA ($Ni^{2+}$). For the liposome permeabilization assay at low BakΔTM concentrations (200 nM), 5% 18:1 DGS-NTA ($Ni^{2+}$) was used, and the mass of PC was adjusted to compensate. The lipids (5 mg) were mixed in chloroform and dried under nitrogen gas flow, resuspended in 0.25 mL of assay buffer (10 mM HEPES pH 7.0, 200 mM KCl, 5 mM $MgCl_2$), sonicated in a sonication bath until homogenous, subjected to five freeze-thaw cycles, then extruded using a 100 nm polycarbonate membrane. The liposomes were prepared with the addition of the polyanionic dye 8-aminoaphthalene-1,3,6-trisulfonic acid (ANTS) and the cationic quencher *p*-xylene-bis-pyridinium bromide (DPX) as described [14,15]. For experiments at low BakΔTM concentrations (200 nM), 0.05 mg/mL OMM-like lipids doped with 5% (w/w) 18:1 DGS-NTA ($Ni^{2+}$) were used. For experiments under BakΔTM auto-active conditions, 0.3 mg/mL OMM-like lipids doped with 10% (w/w) 18:1 DGS-NTA ($Ni^{2+}$) were used. All resulting data were averaged from three technical replicates unless a clear outlier due to experimental artifact was removed. Data analysis was performed by first normalizing to the minimum value in each dataset, then calculating the mean for the technical replicates. The data represented in each of the figures was scaled such that 0 represents the BakΔTM-BclxLΔTM complex and 1 represents either auto-activated BakΔTM or cBid-activated BakΔTM. The data are represented as relative ANTS fluorescence (%).

**Peptide synthesis**

The peptides were prepared according to the standard Fmoc-Solid Phase Peptide Synthesis (Fmoc-SPPS) using a tritylchloride polystyrene (TCP) resin[16]. The completion of the coupling reactions was controlled by analytical ESI-MS. If necessary, the coupling reactions were repeated, especially when Fmoc-Asp(O$^t$Bu)-OH was coupled to the H-Leu-rest. Following acid labile groups were used for protection of side-chains of amino acids: Pbf for Arginine; *t*Bu for Threonine, Serine, Aspartic acid, and Tyrosine; Boc for Lysine. They were removed using a mixture of trifluoroacetic acid (TFA) / DCM / triisopropylsilane (TIPS) / water (80:10:5:5) for 1.5 hours at RT. The resulting peptides were purified by semi-preparative HPLC and their purity was confirmed by analytical HPLC-ESI-MS.

**HPLC and mass spectrometry**

Analytical HPLC-ESI-MS was performed on a Hewlett-Packard Series HP 1100 equipped with a Finnigan LCQ mass spectrometer using a YMC-Hydrosphere C18 column (12 nm pore size,

3 μm particle size, 125 mm×2.1 mm) or YMC-Octyl C8 column (20 nm pore size, 5 μm particle size, 250 mm×2.1 mm) and $H_2O$ (0.1% *v/v* formic acid) / MeCN (0.1% *v/v* formic acid) as eluents. Semi-preparative HPLC was performed using a Beckmann instrument (system gold, solvent delivery module 126, UV detector 166), an YMC ODS-A column (20×250 mm, 5 μm), flow rate: 8 mL/min, linear gradients of $H_2O$ (0.1% *v/v* TFA) and MeCN (0.1% *v/v* TFA).

**Circular dichroism (CD) spectroscopy**

CD spectra and thermal transitions were measured with a Jasco J-1500 spectropolarimeter. Secondary structure estimation based on CD spectra was done with the BestSel server[17]. Thermal melting experiments were conducted at a heating rate of 1°C/min and at a wavelength of 220 nm (monitoring the α-helical content of the protein) and analyzed with a Boltzmann equation[18].

**Isothermal titration calorimetry (ITC)**

ITC experiments to characterize the affinity between BclxL and VDAC1 peptides were conducted with a MicroCal PEAQ-ITC (Malvern Panalytical) at 20°C. Typically 20 μM of BclxL in 10 mM NaPi pH 7.0, 50 mM NaCl, 0.5 mM EDTA and 1 mM DTT in the cell was titrated with 200 μM VDAC1 peptide (linear and stapled) in the syringe using 20 2 μL injections. Data were analyzed by the MicroCal PEAQ-ITC Analysis Software v1.41.

**Crystallization, diffraction data collection and processing**

The crystallization experiments with the single-chain BclxLΔLT-VDAC1-N construct were performed at the X-ray Crystallography Platform at Helmholtz Munich. The initial crystallization screening was done at 292 K using 10 mg/mL of protein with a nanodrop dispenser in sitting-drop 96-well plates and commercial screens. After selecting the best hits from the screening, manual optimization was performed. The best X-ray diffraction data set was collected for a crystal grown in 1.6 M ammonium sulfate, 0.1 M bicine pH 9.0. For the X-ray diffraction experiments, the crystals were mounted in a nylon fiber loop and flash-cooled to 100 K in liquid nitrogen. The cryoprotection was performed for 2 seconds in reservoir solution complemented with 20% (v/v) ethylene glycol. Diffraction data was collected on the X06SA beamline (SLS, Villigen, Switzerland) at 100 K. Data set was indexed and integrated using XDS[19] (v. 01.10.14) and scaled using SCALA[20,21] (v. 01.10.14). Intensities were converted to structure-factor amplitudes using the program TRUNCATE[22] (v. 8.0.019). Table S1 summarizes data collection and processing statistics.

**X-ray structure determination and refinement**

The structure of BclxL-VDAC1 complex was solved by molecular replacement (PHASER[23], v. 2.8.3) using the structure of BclxL published by Muchmore *et al.*, (PDB: 1MAZ[24]). Model rebuilding was performed in COOT[25] (v. 0.9.8.93). The refinement was done in REFMAC5[26] (v. 5.8.0158) using the maximum-likelihood target function. The stereochemical analysis of the final model was done in PROCHECK[27] (v. 3.5.4.) and MolProbity[28] (v. 4.02b-467). The final model is characterized by $R_{work}$ and $R_{free}$ factors of 16.98% and 20.06%, respectively (Table S1). Atomic coordinates and structure factors have been deposited in the Protein Data Bank under accession code 9HPS.

**NMR spectroscopy**

NMR experiments were recorded on Bruker spectrometers operating at 500 to 950 MHz proton frequency equipped with cryogenic probes. Spectral analysis was done with NMRFAM-SPARKY [29].

NMR titrations of *U*-[$^{2}$H, $^{15}$N]- ε-$^{13}CH_3$-methionine-labeled VDAC1-V3M in nanodiscs with BclxLΔTM (residues 1-213) were conducted on an 800 MHz $^{1}$H frequency spectrometer. Experiments were conducted in 20 mM NaPi pH 6.8, 50 mM NaCl, 0.5 mM EDTA and 2 mM DTT with 7% $D_2O$ at 313 K. 2D-[$^{13}$C, $^{1}$H]-HMQC experiments with VDAC1-V3M in MSPΔH5 (50 μM) and VDAC1-V3M in MSP1D1 (75 μM) were recorded with 64 transients per increment and 48 complex data points in the indirect $^{13}$C dimension. BclxLΔTM was then added stepwise at concentrations of 100 μM, 500 μM and 1 mM.

NMR titrations of $^{2}$H, $^{15}$N-labeled BclxLΔTM with VDAC1 in MSPΔH5 nanodiscs (8 nm diameter) were conducted at 950 MHz proton frequency. A 2D-[$^{15}$N,$^{1}$H]-TROSY spectrum of 100 μM $^{2}$H, $^{15}$N-labeled BclxLΔTM in 20 mM NaPi pH 6.5, 50 mM NaCl, 0.5 mM EDTA and 2 mM DTT with 7% $D_2O$, supplemented with protease inhibitor (cOmplete™, Roche) was recorded at 303 K. Subsequently, VDAC1 nanodiscs were added to at molar ratios of 1:1, 1:2, 1:4 and 1:10. The 1:2 titration point (100 μM $^{2}$H, $^{15}$N-labeled BclxLΔTM + 200 μM VDAC1 in MSPΔH5 nanodiscs) was used for analysis. To exclude effects of the interaction of BclxLΔTM with the lipid bilayer surface of the nanodisc particle, a 2D-[$^{15}$N,$^{1}$H]-TROSY spectrum of 100 μM $^{2}$H,$^{15}$N-labeled BclxLΔTM in presence of 200 μM empty MSPΔH5 nanodiscs was recorded and used as a reference. An identical workflow was done for probing the interaction between $^{2}H$, $^{15}N$-labeled BclxLΔTM and VDAC1 in MSP1D1 lipid nanodiscs (10 nm diameter). Furthermore, 100 µM $^{2}H$, $^{15}N$-labeled BclxLΔTM in complex with 100 µM PUMA BH3 peptide was titrated with 200 µM VDAC1 in MSPΔH5 nanodiscs and compared to the 2D-[$^{15}N$,$^{1}H$]-TROSY spectrum of $^{2}H$, $^{15}N$-labeled BclxLΔTM in complex with the PUMA BH3 peptide.

The interaction between $^{2}H$,$^{15}N$-labeled BclxLΔLT (residues 1-44 and 85-213) and the linear and stapled VDAC1-N peptides (residues 1-24 and 7-21 stapled between residues 11 and 15) was probed by NMR 2D-[$^{15}N$,$^{1}H$]-TROSY experiments at 303K and at 500 or 800 MHz 1H frequency, respectively. Buffer was 20 mM NaPi, pH 7.0, 50 mM NaCl, 0.5 mM EDTA, 5 mM DTT. We used 100 µM of $^{2}H$,$^{15}N$-labeled BclxLΔLT and added the respective peptide until no marked chemical shift perturbation changes could be observed (10-fold exess for the linear peptide and 2.5-fold excess for the stapled peptide). The obtained binding curves were analyzed by a one-site binding model. For the stapled peptide, binding was also monitored with 100 µM *U*-$^{2}H$,$^{15}N$, Ile-$\delta_1$,Leu-$\delta_2$, Val-$\gamma_2$, Ala-β-[$^{13}CH_3$]-labeled BclxLΔLT using 2D-[$^{13}C$,$^{1}H$]-HMQC experiments measured at 303K and at 800 MHz $^{1}H$ frequency.

NMR was also used to obtain binding affinities of VDAC1-N peptides in an alanine scanning experiment. VDAC1-N peptides (Table S2) were titrated into 100 µM $^{2}H$,$^{15}N$-labeled BclxLΔLT and the resulting chemical shift perturbation was plotted against the peptide concentration to extract $K_D$ values for each peptide using a one-site binding model.

NMR spectra of 100 µM $^{1}H$,$^{15}N$-labeled VDAC1-N peptide were obtained at 303K at 500 MHz $^{1}H$ frequency in 20 mM NaPi, pH 7.0, 50 mM NaCl, 0.5 mM EDTA, 5 mM DTT. For mapping the binding site within the peptide for BclxL, a 5-fold excess of BclxLΔTM was added and chemical shift perturbations were mapped on the structure of VDAC1-N extracted from full-length VDAC1[1].

**Cryo-EM sample preparation**

For cryo-EM sample preparation, 4 µL protein solution prepared as mentioned above was applied to a glow-discharged UltrAuFoil R 1.2/1.3, 300 mesh (Quantifoil Micro Tools). The grids were blotted for 4.5 s and plunge-frozen in liquid ethane after 1 s of drain time using a blot force of -1, in 100% relative humidity at 13°C (Vitrobot Mark IV, Thermo Fischer Scientific). The plunged EM grids were clipped and used for automated dataset collection using the EPU software with a Krios G4 Cryo-TEM (Thermo Fischer Scientific) at 300 kV, equipped with an E-CFEG, a Selectris X Energy Filter at a slit width of 10 eV, and Falcon 4i Direct Electron Detector.

**Cryo-EM data processing**

Movies for all datasets were monitored for quality and preprocessed on CryoSPARC Live[30,31]. The gain-corrected, motion-corrected, dose-weighted, and CTF-estimated micrographs were exported for further processing to CryoSPARC (v3.1.0-v4.5.1). For all subsequent jobs described, the following modifications were made to the default settings:
*2D Classification*: class2D_max_res and class2D_max_res_align were usually increased to 9 and 12 respectively, class2D_force_max was set to false, class2D_num_full_iter was set to 2-4, class2D_num_full_iter_batch was set to 40-100, and class2D_num_full_iter_batchsize_per_class was set to 400-1000.
*Ab-Initio Reconstruction*: abinit_max_res was decreased to 6-9, abinit_init_res was reduced to 15-25, abinit_minisize_init was set to 300, abinit_minisize was set to 1000.
*Non-Uniform Refinement*: refine_res_init was usually set to 6-12, and refine_defocus_refine and refine_ctf_global_refine were set to true in all datasets but the nanodisc one. refine_do_ews_correct was set to true, given prior knowledge of our microscope.
A compilation of jobs created and the processing tree can be seen in Fig. S4. The data acquisition statistics can be seen in Tab. S3.

**Molecular dynamics (MD) simulations**

The complex between VDAC1 and the soluble domain of BclxL was assembeled using the herein determined crystal structure of BclxL in complex with VDAC1-N. The C-terminal end of VDAC1-N in the complex was linked to the β-barrel of VDAC1 using Chimera[32]. This assembled complex between VDAC1 and BclxL was inserted into a hexagonal box of 1,2-di-myristoyl-sn-glycero-3-phosphocholine (DMPC) and 1,2-di-myristoyl-sn-glycero-3-phosphoglycerol (DMPG) (3:1 ratio) lipid bilayer using the CHARMM-GUI web server (http:/www.charmm-gui.org)[33,34] in water supplemented with 0.15 M KCl. Equilibration of the system was done at 310 K in two phases with 3 cycles each. The force constants to fix the position of the protein and membrane were gradually reduced in each cycle. In the first phase, a timestep of 1 fs and a simulation time of 50 ps in each cycle were used whereas in the second phase a timestep of 2 fs and a simulation time of 200 ps for each cycle were used. Total equilibration time was 750 ps. The production MD simulation of 100 ns duration was carried out with the isothermal-isobaric ensembles at 310 K with the program NAMD[35] in the absence of a transmembrane potential. A smoothing function was applied to truncate short-range electrostatic interactions. Visualization of the final structural model was done with VMD[36]. VDAC1 with its N-terminal helix located in its canonical internal position as well as in an outside position, as observed in the complex with BclxL was also subjected to an extended MD simulation in a POPC:POPG (3:1) lipid bilayer environment. The simulation was set up as described above at 310K but the simulations were run for 5 μs using the CUDA-enhanced version of GROMACS[37] on an in-house GPU workstation.

**Supporting Information Tables**

**Table S1: X-ray data collection and refinement statistics for BclxL in complex with VDAC1-N.**

| **Data collection** | |
|---|---|
| PDB ID | 9HPS |
| Beamline | SLS X06SA PX |
| Wavelength | 0.99999 |
| Space group | $C222_1$ |
| Cell dimensions *a*, *b*, *c* (Å) | 35.12, 98.75, 103.91 |
| Resolution (Å) | 50–1.95 (2.00–1.95)* |
| $R_{merge}$ | 10.0 (64.6) |
| $I / \sigma I$ | 12.6 (2.3) |
| CC (1/2) | 99.7 (80.2) |
| Completeness (%) | 98.7 (96.2) |
| Redundancy | 4.1 (3.5) |
| **Refinement** | |
| Resolution (Å) | 1.95 |
| No. reflections | 12,735 |
| $R_{work}$ / $R_{free}$ | 16.98 / 20.06 |
| No. atoms | |
| Protein | 1,206 |
| Peptide | 136 |
| Water | 80 |
| Other | 15 |
| *B*-factor overall | 36.1 |
| R.m.s. deviations | |
| Bond lengths (Å) | 0.008 |
| Bond angles (°) | 1.76 |
| Ramachandran plot | |
| Most favored (%) | 96 |
| Additional allowed (%) | 4 |

*Values in parentheses are for highest-resolution shell.

**Table S2: Alanine scan peptides derived from VDAC1-N.**

| Sequence (Y7 to G21) | *m*/*z*, [M+H$^+$] (calc) | *m*/*z*, [M+H$^+$] (found) | Purity (UV) |
|---|---|---|---|
| Y A D L G K S A R D V F T K G | 1627.8 | 1627.9 | 97% |
| A A D L G K S A R D V F T K G | 1536.8 | 1536.8 | >98% |
| Y A A L G K S A R D V F T K G | 1583.7 | 1583.9 | >98% |
| Y A D A G K S A R D V F T K G | 1585.8 | 1585.6 | >98% |
| Y A D L A K S A R D V F T K G | 1641.9 | 1641.8 | >98% |
| Y A D L G A S A R D V F T K G | 1570.8 | 1571.1 | 95% |
| Y A D L G K A A R D V F T K G | 1611.8 | 1611.7 | >98% |
| Y A D L G K S A A D V F T K G | 1542.8 | 1542.7 | 95% |
| Y A D L G K S A R A V F T K G | 1583.9 | 1583.9 | >98% |
| Y A D L G K S A R D A F T K G | 1599.8 | 1599.7 | 98% |
| Y A D L G K S A R D V A T K G | 1551.8 | 1551.9 | 96% |
| Y A D L G K S A R D V F A K G | 1597.8 | 1597.7 | 98% |

**Table S3: Cryo-EM data acquisition statistics.**

| Sample | cMSP1D1-VDAC1 monomer | cMSP1D1-VDAC1 dimer | cMSP1ΔH5-VDAC1 |
|---|---|---|---|
| **Data collection and processing** | | | |
| Magnification | 270,000 | 270,000 | 270,000 |
| Voltage (kV) | 300 | 300 | 300 |
| Electron exposure ($e^-$/Å$^2$) | 60 | 60 | 50 |
| Defocus range (μm) | 0.8-2.0 | 0.8-2.0 | 0.4-2.0 |
| Pixel size (Å) | 0.46 | 0.46 | 0.46 |
| Symmetry imposed | C1 | C1 | C1 |
| Movies (No.) | 10,258 | 12,000 | 20,380 |
| Initial particle images (No.) | 1,800,035 | 179,411 | 4,011,760 |
| Final particle images (No.) | 169,625 | 31,802 | 245,265 |
| Map resolution (Å) | 7.21 | 6.99 | 5.70 |
| Map resolution range (Å) | 2.50-7.80 | 3.19-6.72 | 5.45-6.46 |
| FSC threshold | 0.143 | 0.143 | 0.143 |

### Supporting Information Figures

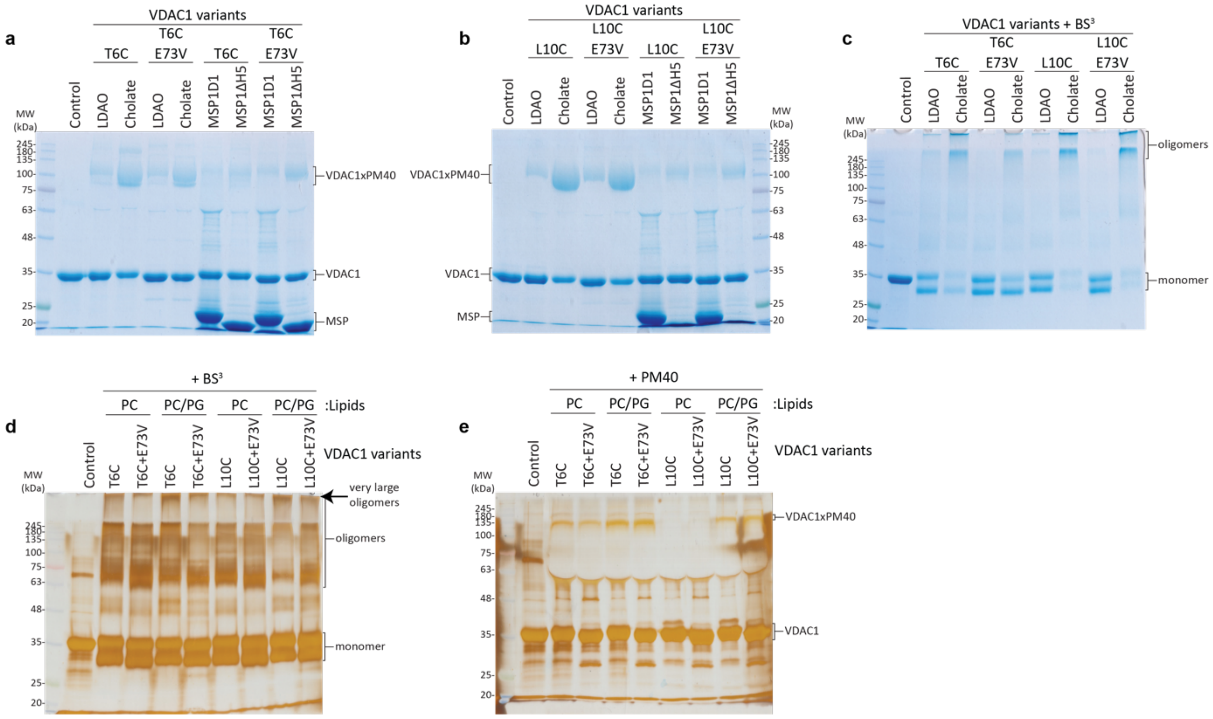

**Figure S1. Chemical crosslinking and modification to probe VDAC1 oligomerization and α-helix exposure.** (a) Chemical modification of VDAC1-T6C with polyethyleneglycol (PEG)-maleimide of 40 kDa (PM40) in different detergents and lipid nanodiscs. The VDAC1 point mutation E73V, that was reported to show a tighter interaction between the α-helix and the β-barrel wall [38], was also assayed. (b) Same as in (a) but with the VDAC1 variant L10C that is located further inside the pore. (c) Chemical crosslinking of VDAC1 T6C or L10C variants with the amino-selective crosslinker bissulfosuccinimidyl suberate ($BS^3$). (d) $BS^3$ crosslinking of VDAC1 in liposomes composed of POPC or POPC/POPG (75%:25%) lipids. Very large VDAC1 oligomers can only be observed in the presence of a negatively charged lipid surface. (e) PM40 modification of the samples shown in (d). The negatively charged lipid POPG was required for VDAC1 α-helix exposure which can be best seen with the L10C variant.

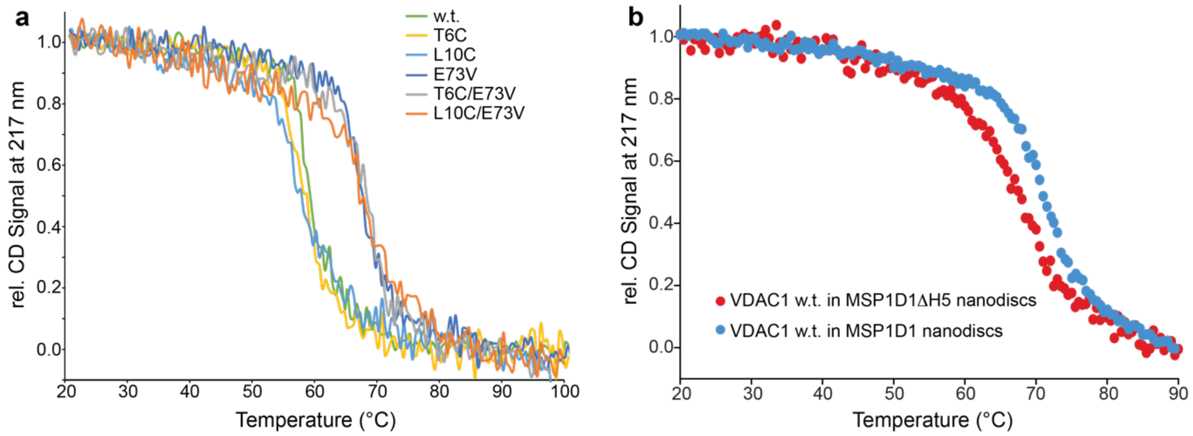

**Figure S2. Thermal stabilities of VDAC1 variants in detergent micelles and nanodiscs.** CD-detected thermal melting experiments conducted in a 1 mm pathlength cuvette. The signal was normalized using the CD signal intensity at 20°C and 100 or 90°C, respectively. The heating rate was 1°C/min. (a) VDAC1 variants as indicated in the plot in 0.1% LDAO, 20 mM NaPi pH 7.0, 1 mM DTT. (b) VDAC1 wild-type in MSP1D1 (blue) and MSP1D1ΔH5 (red) nanodiscs loaded with DMPC:DMPG=3:1 lipids in the same buffer.

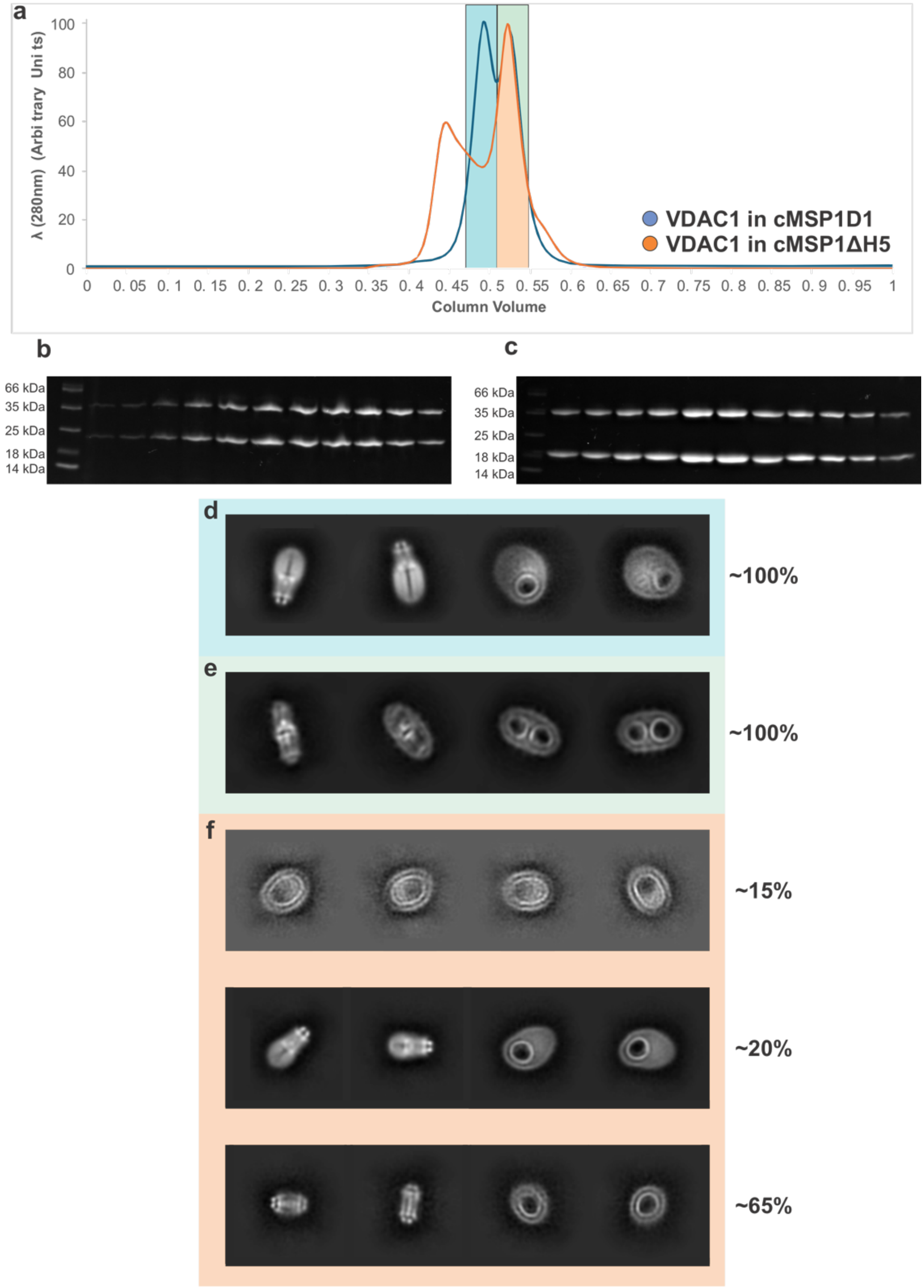

**Figure S3. Homogeneity of VDAC1 nanodisc samples and cryo-EM analysis**
(**a**) Comparative size exclusion chromatogram of both samples. Fractions collected are highlighted. (**b**) SDS-PAGE of cMSP1D1 VDAC1 (**c**) SDS-PAGE of cMSP1ΔH5 VDAC1(**d-f**) Representative 2D class averages of species found in each sample. (**d**) Large nanodiscs containing a single VDAC1 protomer from the early peak of cMSP1D1 VDAC1 nanodiscs. (**e**) Large nanodiscs containing a single VDAC1 dimer from the early peak of cMSP1D1 VDAC1 nanodiscs. (**f**) Variety of species found in the cMSP1ΔH5 VDAC1 dataset and their relative abundance. Larger cMSP1ΔH5 nanodiscs are most likely formed by dimeric cMSP that co-purifies with monomeric cMSP[39].

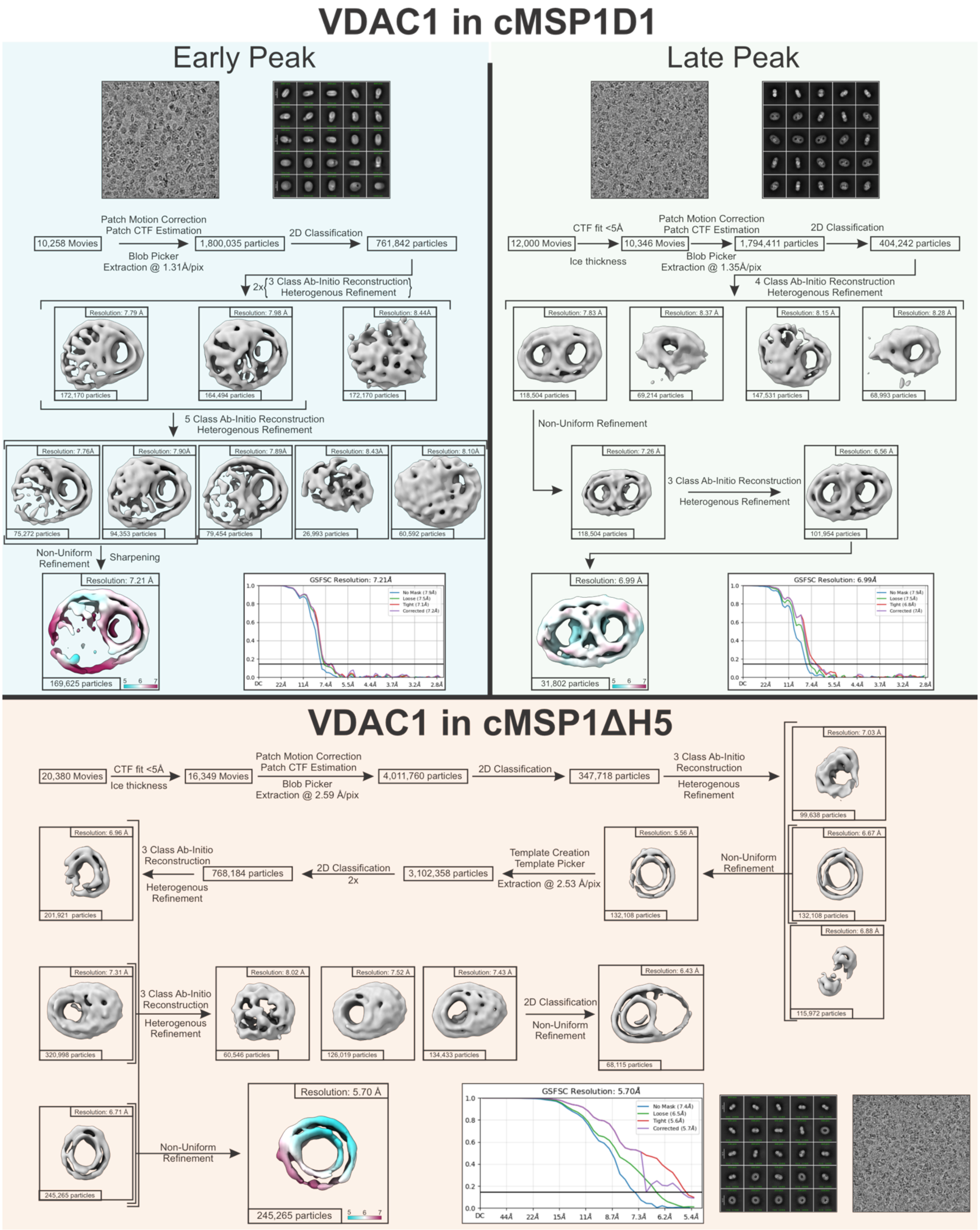

**Figure S4. Cryo-EM processing workflow of VDAC1 in nanodiscs of different sizes.**
**Top left**: Processing of cMSP1D1 VDAC1 monomers in nanodiscs (peak 1) yielding a final resolution of 7.21 Å. **Top Right**: Processing of cMSP1D1 VDAC1 dimers in nanodiscs (peak 2) giving a final resolution of 6.99 Å. **Bottom**: Processing of cMSP1ΔH5 VDAC1 nanodiscs yielding a resolution of 5.7 Å.

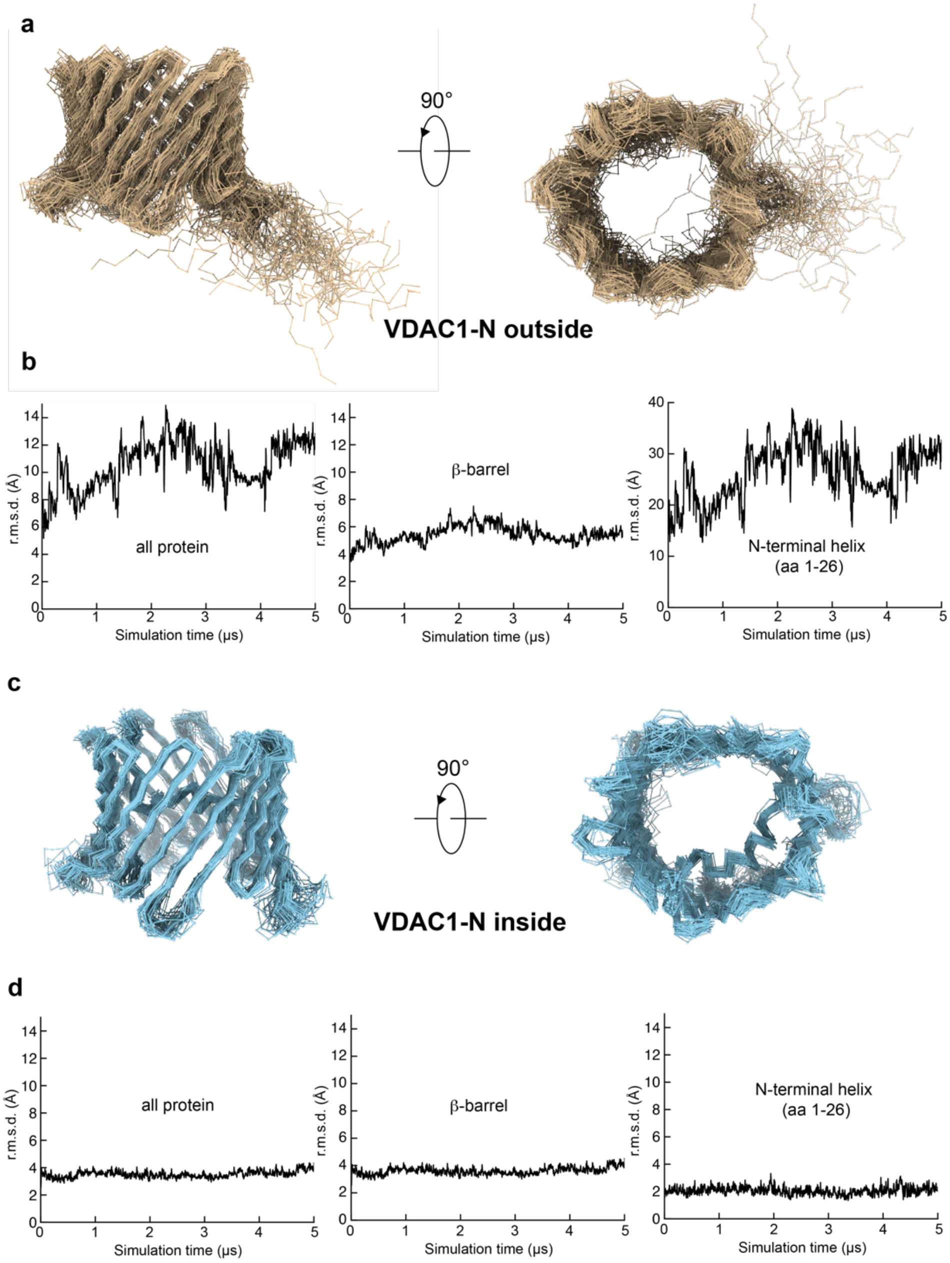

**Figure S5. Molecular dynamics (MD) simulations of VDAC1 with the N-terminal helix outside or inside the β-barrel.** (a) Overlay of structural snapshots (every 100 ns) along a 5 μs trajectory of VDAC1 with the N-terminal helix outside the β-barrel in a lipid bilayer composed of POPC and POPG (3:1) at T=310 K. (b) The root mean square deviation (r.m.s.d.) of the atomic coordinates of the entire protein, the β-barrel or just the N-terminal helix obtained from the MD trajectory show that VDAC1 undergoes large conformational fluctuations with an almost unrestricted N-terminal helix (r.m.s.d. up to 40 Å) but also with a less well defined β-barrel. (c) Same as in (a) but using the canonical human VDAC1 structure (PDB ID: 5XDO) with the N-terminal helix attached to the inside of the β-barrel. (d) The r.m.s.d. values for the latter simualtion are much lower with a maximum of 4 Å and an even lower value for the N-terminal helix (~2 Å), suggesting that the N-terminal helix leads to a marked stabilization of the overal VDAC1 structure.

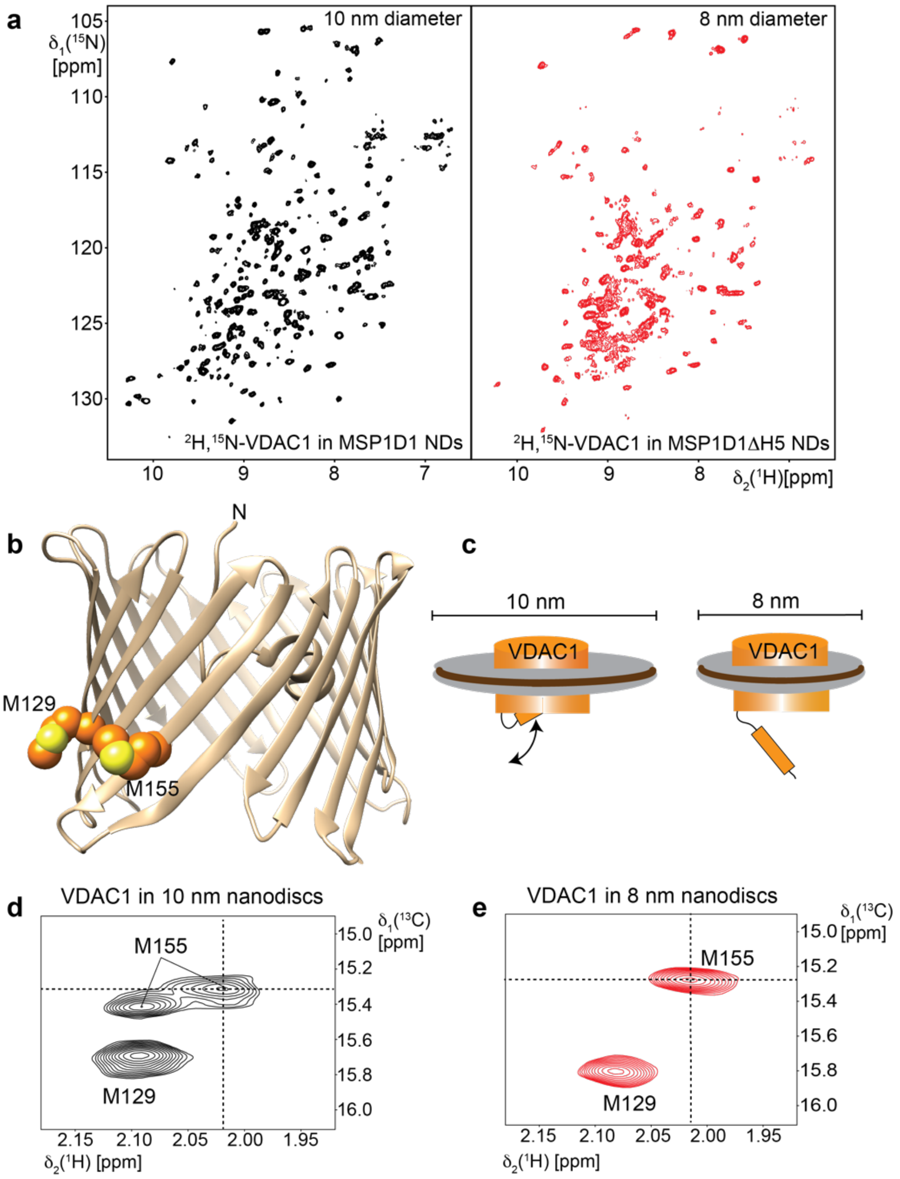

**Figure S6. 2D-NMR spectra of VDAC1 in nanodiscs of different sizes.** (a, left) 2D-[$^{15}$N,$^{1}$H]-TROSY spectrum of 200 µM $^{2}$H,$^{15}$N-labeled VDAC1 in MSP1D1 lipid nanodiscs of 10 nm diameter assembeled with a DMPC:DMPG=3:1 lipid blend, measured at 800 MHz $^{1}$H frequency and at T = 45°C in 20 mM NaPi pH 7.0, 50 mM NaCl, 0.5 mM EDTA, 5 mM DTT buffer. (a, right) 2D-[$^{15}$N,$^{1}$H]-TROSY spectrum of 200 µM $^{2}$H,$^{15}$N-labeled VDAC1 in MSP1D1ΔH5 lipid nanodiscs of 8 nm diameter. The smaller nanodisc leads to marked line broadening of VDAC1 resonances. (b) Structure of VDAC1 with the native methionine residues (M129, M155) labeled. The N-terminal methionine residue is removed during protein expression and not visible in the spectrum. (c) Cartoon presentation of VDAC1 in 8 and 10 nm nanodiscs, with the N-terminus being exposed in the smaller nanodiscs. (d) 2D-[$^{13}$C,$^{1}$H]-HMQC spectrum of Met-ε-$^{13}$C-labeled VDAC1 in 10 nm nanodiscs. As evident from the peak doubling for M155, VDAC1 exists in two conformational states. (e) same as (d) but with VDAC1 in 8 nm nanodiscs. Only one NMR signal is existing for M155 indicating a single conformation. A comparison of the chemical shifts of M155 in both samples, suggests that M155 in 8 nm nanodiscs is present in an exposed conformation and in 10 nm the majority is present in a different β-barrel state.

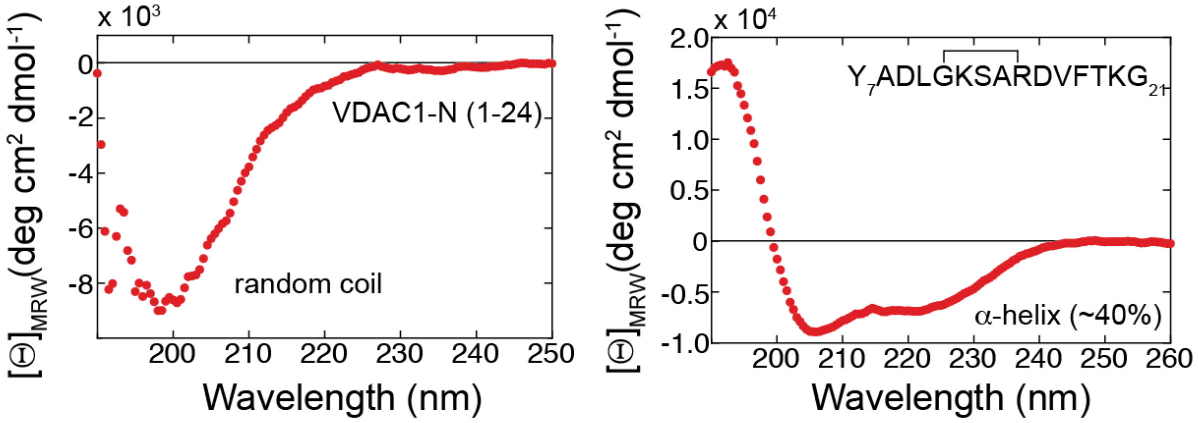

**Figure S7. CD spectra of VDAC1-N and st-VDAC1-N.** (Left) CD spectrum of a peptide derived from the VDAC1 N-terminal segment (residues 1-24) indicates a random coil secondary structure. (Right) The CD spectrum of a hydrocarbon stapled VDAC1 peptide (residues 7-21 with the staple between residues 11 and 15) adopts ~40% α-helical secondary structure, as estimated with the program BestSel[17]. Data are represented as mean residue weight (MRW) ellipticity. A 1 mm path length cuvette was used at T = 20°C.

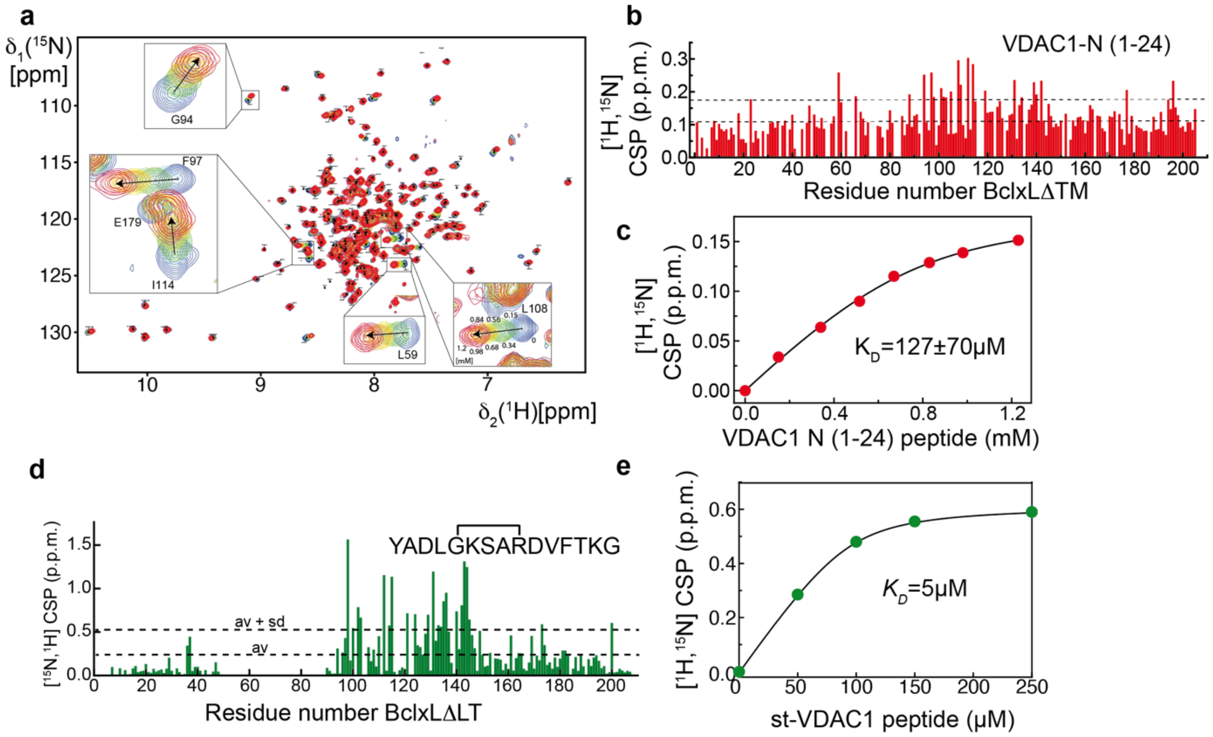

**Figure S8. NMR validation of the interaction between VDAC1-N-terminal peptides and BclxL.** (a) 2D-[$^{15}$N,$^{1}$H]-TROSY NMR titration with $^{2}$H,$^{15}$N-labeled BclxLΔTM (lacking the transmembrane helix) and the linear VDAC1-N-terminal peptide (residues 1-24). Some affected resonances are shown in the insets. (b) Chemical shift perturbation (CSP) values at the highest concentration of VDAC1-N (1.2 mM) for each residue in BclxLΔTM. (c) CSP values in BclxL plotted against the concentration of VDAC1-N, giving rise to a binding isotherm, yielding a $K_D$ value of ~130 µM. (d) CSP pattern of BclxLΔLT (lacking the flexible loop and the transmembrane helix) at a saturating concentration of st-VDAC1-N. The detected CSP amplitude is markedly larger than with the VDAC1-N peptide. (e) Binding isotherm obtained with st-VDAC1-N results in a $K_D$ value of ~5 µM.

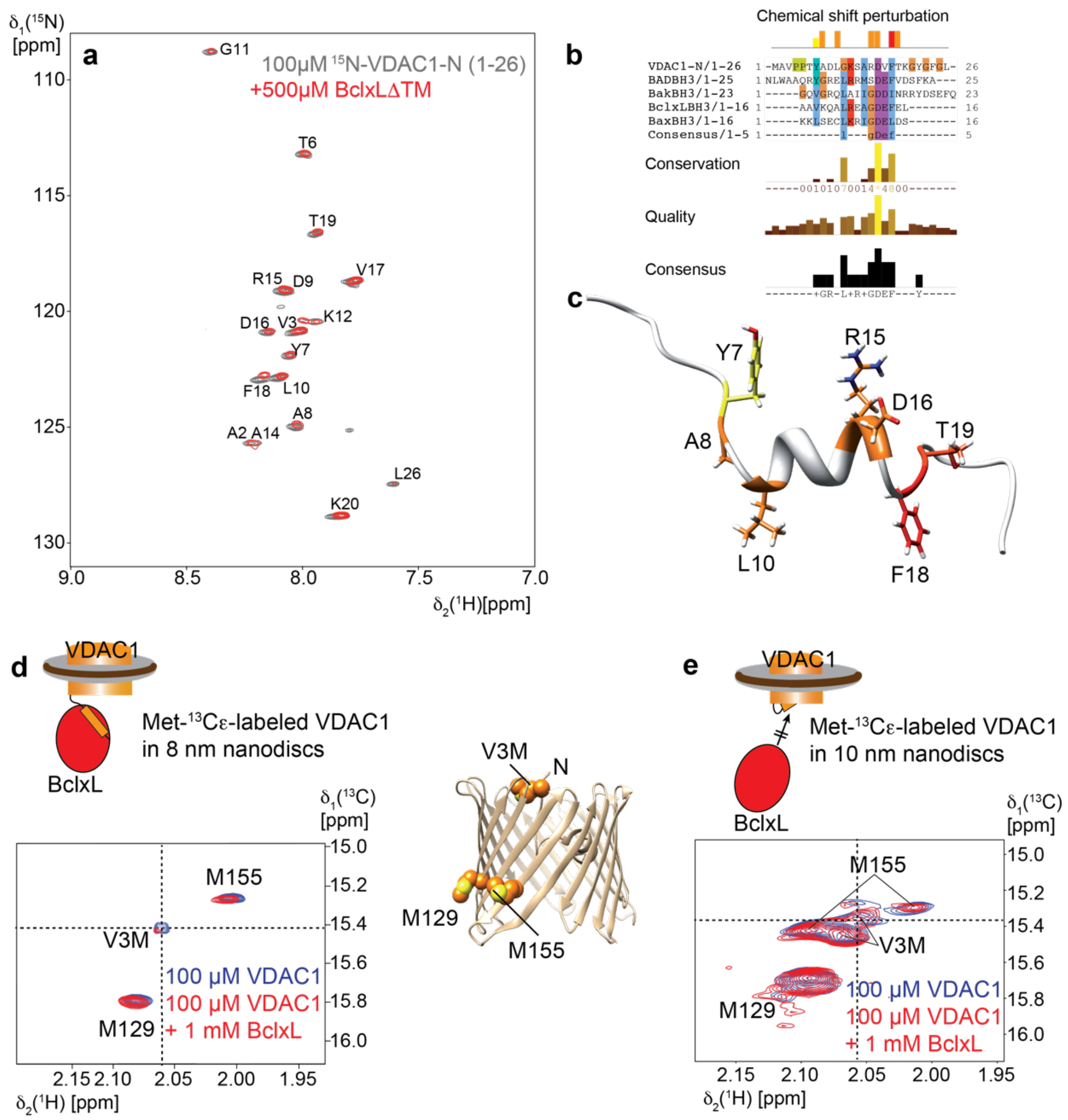

**Figure S9. Interaction between VDAC1-N and BclxL at a per-residue resolution obtained by NMR.** (a) 2D-[$^{15}$N,$^{1}$H]-TROSY spectra of $^{15}$N-labeled VDAC1-N (1-26) [8] and after the addition of a 5-fold amount of BclxL. (b) Chemical shift perturbations (CSPs) were observed for the indicated positions with yellow, orange and red representing weak, medium and strong effects, respectively. Below the CSPs is a multiple seuquence alignemnt of VDAC1-N with BH3 peptides from various Bcl2 proteins, indicating a low level of conservation. (c) CSPs mapped onto the structure of VDAC1-N taken from the NMR structure of full-length VDAC1 in detergent micelles[1]. (d) 2D-[$^{13}$C,$^{1}$H]-HMQC spectrum of Met-ε-$^{13}$C-labeled VDAC1 in 8 nm lipid nanodiscs alone (blue) and in presence of a 10-fold molar excess of BclxL (red). The NMR signal for Met3 is mostly affected by the addition of BclxL. Methionine chemical shift assignments were obtained by mutagenesis. (e) same as in (d) but with VDAC in 10 nm nanodiscs. There is peak doubling for Met3 and Met115, indicative of two conformational states. For Met3, the minor peak (crosshair) is affected by the addition of BclxL. A comarison with the spectrum in (d) suggests that this peak prepresents the exposed conformation of the VDAC1 N-terminus.

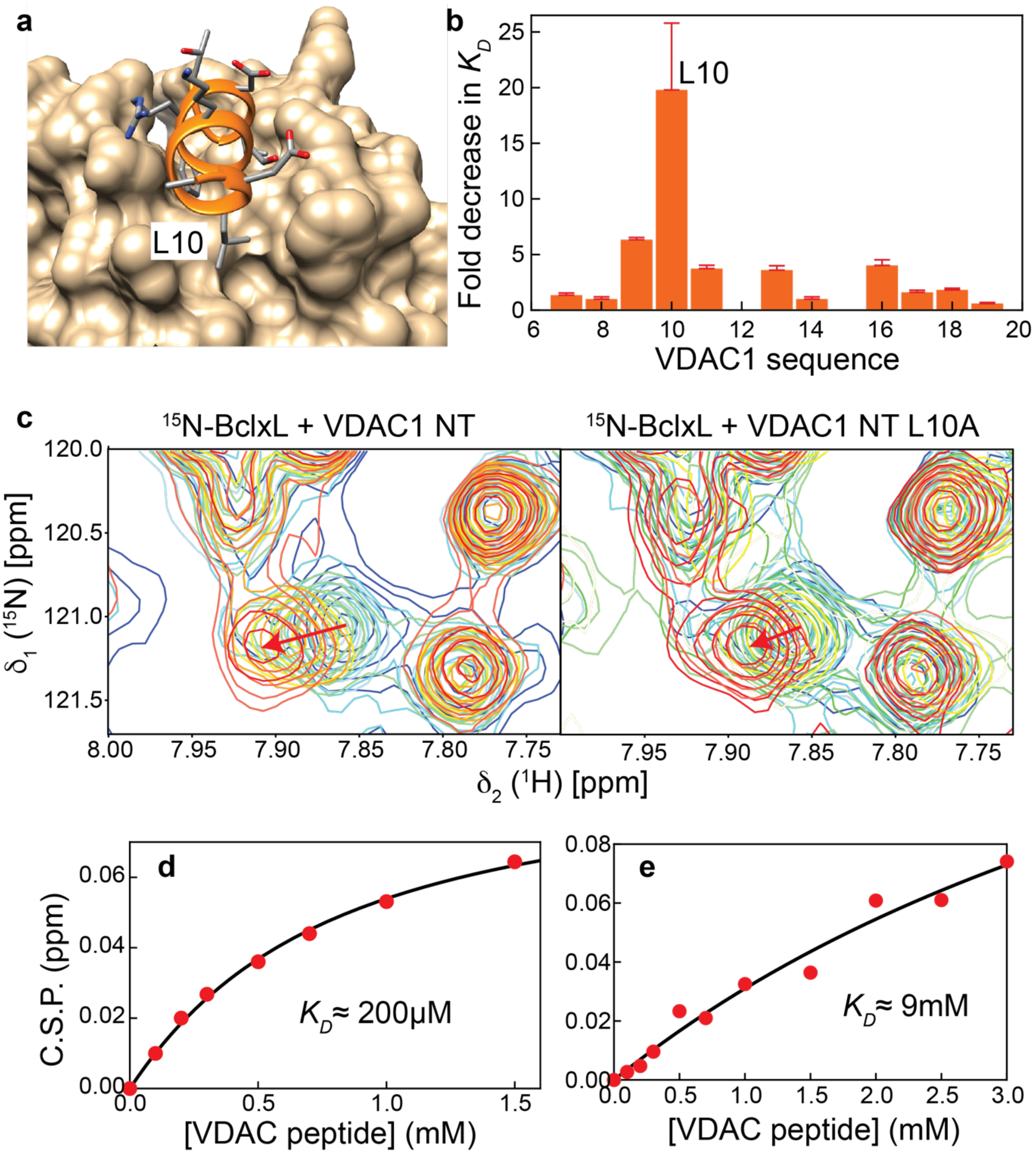

**Figure S10. Alanine scan of the VDAC1-N peptide and its interaction with BclxL.** (a) Structure of BclxL in complex with the VDAC1-N peptide. Sidechains are depicted in a stick representation and leucine 10 (L10) is labelled. (b) Decrease in affinity with the VDAC1-N peptides where the indicated position is replaced by alanine (ala scan). Mutation of Asp9, Leu10, Gly11, Ser13, Asp16 have the strongest impact on the interaction with BclxL. (c) NMR spectral overlays of the titration steps with the wild-type (left) or the L10A peptide (right). (d) NMR-derived binding isotherm with the wild-type VDAC1-N peptide. (e) same as (d) but with the L10A VDAC1-N peptide.

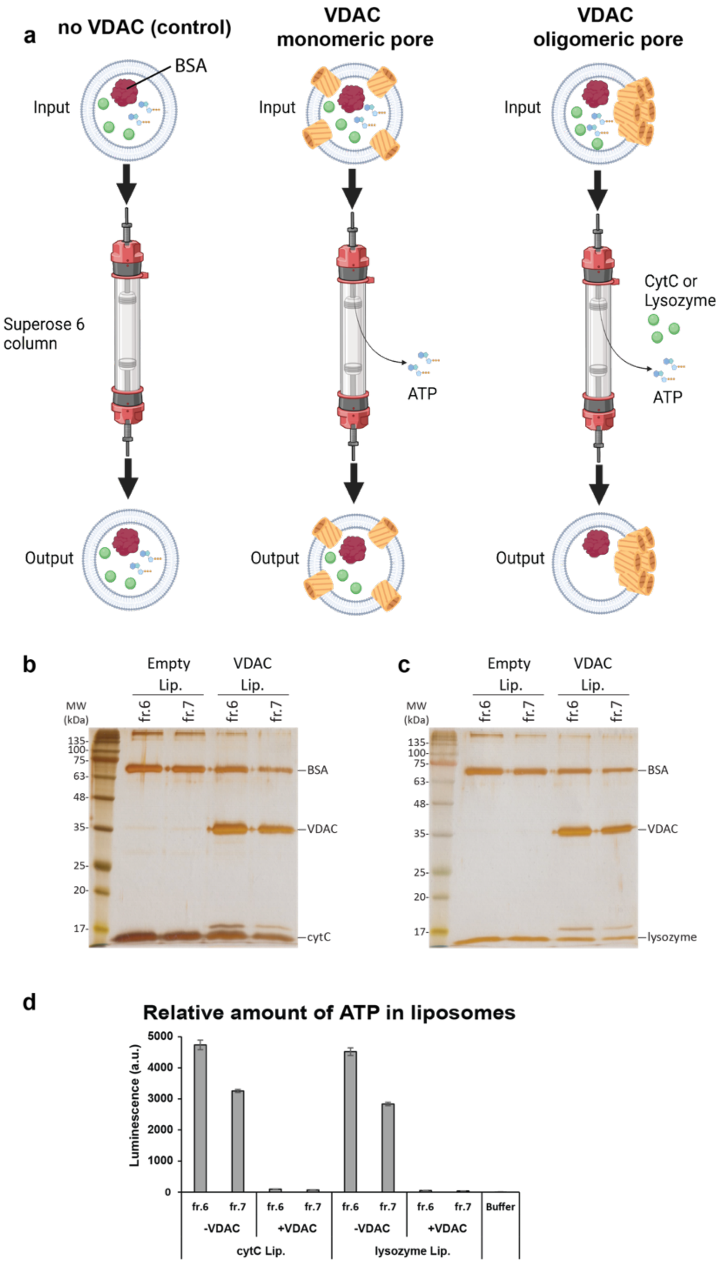

**Figure S11. VDAC1 in liposomes is permeable for ATP but not for small proteins.** (a) Schematic of the size exclusion assay to probe ATP or cytochrome C/lysozyme translocation across VDAC1. (b) Liposomes loaded with bovine serum albumine (BSA) and cytochrome C (cytC) with or without VDAC1 were injected on a Superose 6 size exclusion chromatography column (Cytvia). If proteins inside the liposomes are able exit the liposomes the corresponding band on the SDS-PAGE should be markedly weakened or be completely lost. The ~70 kDa BSA protein serves as a reference since it is assumed to be too large to exit through a VDAC1 pore. For cytC, no clear reduction in the band intensity could be observed. (c) same as in (b) but with lysozyme instead of cytC, showing the same result, i.e. no translocation of the protein across VDAC1. (d) To probe the functionality of VDAC1 and the existence of a pore that is large enough to allow for the transition of metabolites, the amount of ATP in empty and VDAC1-containing liposomes was monitored in the same samples. In contrast to both proteins in (a) and (b) ATP was completely removed in VDAC1 proteoliposomes but not in liposomes without VDAC1, indicating its efficient translocation across VDAC1.

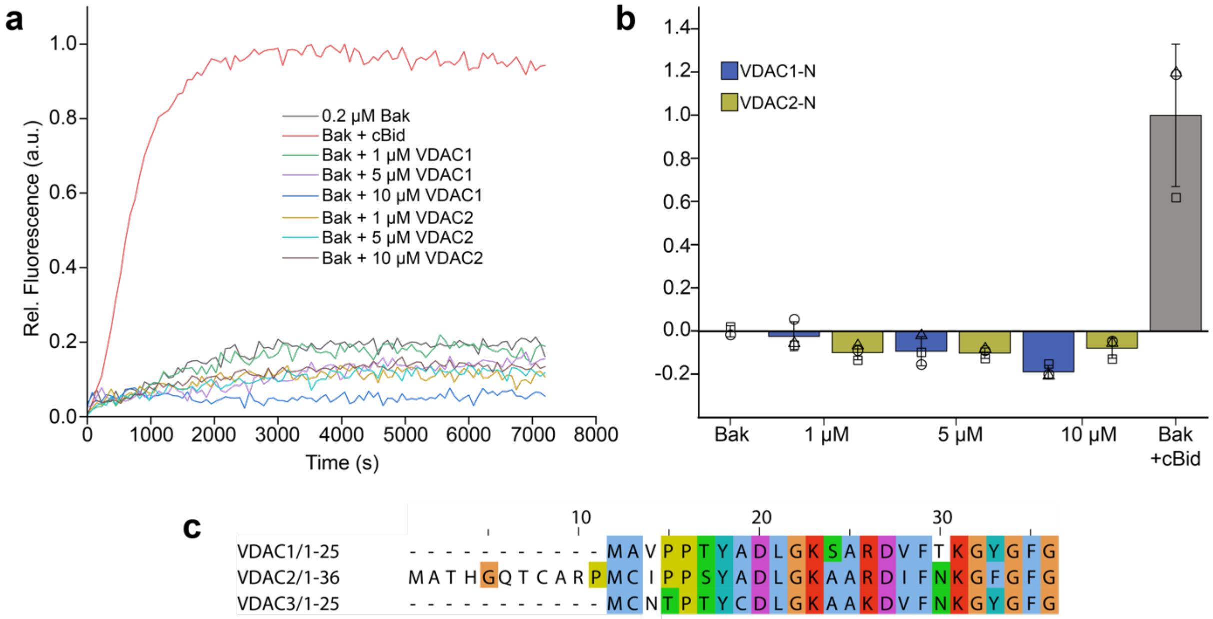

**Figure S12. VDAC-N peptides cannot directly activate pore formation of Bak.** (a) Normalized fluorescence intensity of ANTS in a liposome pore forming assay [14] with 200 nM Bak and different concentrations of VDAC1-N and VDAC2-N peptides, as well as cBid as a positive control. (b) Fluorescence intensity (average fluorescence signal between 4000 and 4500 s) changes from (a) relative to the Bak sample without the additon of peptides or cBid. Bak pore formation at low concentration requires activation by BH3-only proteins, as seen for cBid (50 nM, grey bar). In comparison, up to 10 µM of either GB1-VDAC1-N (blue) or GB1-VDAC2-N (green) peptides were unable to activate BakΔTM. (c) Multiple sequence alignment of human VDAC1, VDAC2 and VDAC3 N-terminal regions shows an almost identical sequence for VDAC1 and VDAC3 and a 10 amino acid residue N-terminal extension in VDAC2. Coloring was done by the Clustral method in Jalview[40].

### Supporting Information References

1 Hiller, S. *et al.* Solution structure of the integral human membrane protein VDAC-1 in detergent micelles. *Science* **321**, 1206-1210 (2008). https://doi.org:10.1126/science.1161302

2 Malia, T. J. & Wagner, G. NMR structural investigation of the mitochondrial outer membrane protein VDAC and its interaction with antiapoptotic Bcl-xL. *Biochemistry* **46**, 514-525 (2007). https://doi.org:10.1021/bi061577h

3 Hausler, E. *et al.* Quantifying the insertion of membrane proteins into lipid bilayer nanodiscs using a fusion protein strategy. *Biochim Biophys Acta Biomembr* **1862**, 183190 (2020). https://doi.org:10.1016/j.bbamem.2020.183190

4 Moldoveanu, T. *et al.* BID-induced structural changes in BAK promote apoptosis. *Nat Struct Mol Biol* **20**, 589-597 (2013). https://doi.org:10.1038/nsmb.2563

5 Raltchev, K., Pipercevic, J. & Hagn, F. Production and Structural Analysis of Membrane-Anchored Proteins in Phospholipid Nanodiscs. *Chemistry* **24**, 5493-5499 (2018). https://doi.org:10.1002/chem.201800812

6 Sperl, L. E., Ruhrnossl, F., Schiller, A., Haslbeck, M. & Hagn, F. High-resolution analysis of the conformational transition of pro-apoptotic Bak at the lipid membrane. *Embo J* **40**, e107159 (2021). https://doi.org:10.15252/embj.2020107159

7 Hagn, F. *et al.* BclxL changes conformation upon binding to wild-type but not mutant p53 DNA binding domain. *J Biol Chem* **285**, 3439-3450 (2010). https://doi.org:10.1074/jbc.M109.065391

8 Reif, M. M., Fischer, M., Fredriksson, K., Hagn, F. & Zacharias, M. The N-Terminal Segment of the Voltage-Dependent Anion Channel: A Possible Membrane-Bound Intermediate in Pore Unbinding. *J Mol Biol* **431**, 223-243 (2019). https://doi.org:10.1016/j.jmb.2018.09.015
9 Ritchie, T. K. *et al.* Chapter 11 - Reconstitution of membrane proteins in phospholipid bilayer nanodiscs. *Methods Enzymol* **464**, 211-231 (2009). https://doi.org:10.1016/S0076-6879(09)64011-8
10 Hagn, F., Etzkorn, M., Raschle, T. & Wagner, G. Optimized phospholipid bilayer nanodiscs facilitate high-resolution structure determination of membrane proteins. *J Am Chem Soc* **135**, 1919-1925 (2013). https://doi.org:10.1021/ja310901f
11 Hagn, F., Nasr, M. L. & Wagner, G. Assembly of phospholipid nanodiscs of controlled size for structural studies of membrane proteins by NMR. *Nat Protoc* **13**, 79-98 (2018). https://doi.org:10.1038/nprot.2017.094
12 Daniilidis, M., Brandl, M. J. & Hagn, F. The Advanced Properties of Circularized MSP Nanodiscs Facilitate High-resolution NMR Studies of Membrane Proteins. *J Mol Biol* **434**, 167861 (2022). https://doi.org:10.1016/j.jmb.2022.167861
13 Gunsel, U. *et al.* Structural basis of metabolite transport by the chloroplast outer envelope channel OEP21. *Nat Struct Mol Biol* **30**, 761-769 (2023). https://doi.org:10.1038/s41594-023-00984-y
14 Kale, J., Chi, X., Leber, B. & Andrews, D. Examining the molecular mechanism of bcl-2 family proteins at membranes by fluorescence spectroscopy. *Methods Enzymol* **544**, 1-23 (2014). https://doi.org:10.1016/B978-0-12-417158-9.00001-7
15 Yethon, J. A., Epand, R. F., Leber, B., Epand, R. M. & Andrews, D. W. Interaction with a membrane surface triggers a reversible conformational change in Bax normally associated with induction of apoptosis. *J Biol Chem* **278**, 48935-48941 (2003). https://doi.org:10.1074/jbc.M306289200
16 Chatterjee, J., Laufer, B. & Kessler, H. Synthesis of N-methylated cyclic peptides. *Nat Protoc* **7**, 432-444 (2012). https://doi.org:10.1038/nprot.2011.450
17 Micsonai, A. *et al.* Accurate secondary structure prediction and fold recognition for circular dichroism spectroscopy. *Proc Natl Acad Sci U S A* **112**, E3095-3103 (2015). https://doi.org:10.1073/pnas.1500851112
18 Privalov, P. L. Stability of proteins: small globular proteins. *Adv Protein Chem* **33**, 167-241 (1979).
19 Kabsch, W. XDS. *Acta Crystallogr D Biol Crystallogr* **66**, 125-132 (2010). https://doi.org:10.1107/s0907444909047337
20 Evans, P. Scaling and assessment of data quality. *Acta Crystallogr D Biol Crystallogr* **62**, 72-82 (2006). https://doi.org:10.1107/s0907444905036693
21 Winn, M. D. *et al.* Overview of the CCP4 suite and current developments. *Acta Crystallogr D Biol Crystallogr* **67**, 235-242 (2011). https://doi.org:10.1107/s0907444910045749
22 French, S. & Wilson, K. On the treatment of negative intensity observations. *Acta Crystallographica Section A* **34**, 517-525 (1978). https://doi.org:doi:10.1107/S0567739478001114
23 McCoy, A. J. *et al.* Phaser crystallographic software. *J Appl Crystallogr* **40**, 658-674 (2007). https://doi.org:10.1107/s0021889807021206
24 Muchmore, S. W. *et al.* X-ray and NMR structure of human Bcl-xL, an inhibitor of programmed cell death. *Nature* **381**, 335-341 (1996). https://doi.org:10.1038/381335a0
25 Emsley, P., Lohkamp, B., Scott, W. G. & Cowtan, K. Features and development of Coot. *Acta Crystallogr D Biol Crystallogr* **66**, 486-501 (2010). https://doi.org:10.1107/s0907444910007493
26 Murshudov, G. N., Vagin, A. A. & Dodson, E. J. Refinement of macromolecular structures by the maximum-likelihood method. *Acta Crystallogr D Biol Crystallogr* **53**, 240-255 (1997). https://doi.org:10.1107/s0907444996012255
27 Laskowski, R. A., Rullmann, J. A. C., MacArthur, M. W., Kaptein, R. & Thornton, J. M. AQUA and PROCHECK-NMR: Programs for checking the quality of protein structures solved by NMR. *J Biomol NMR* **8**, 477-486 (1996).

28 Chen, V. B. *et al.* MolProbity: all-atom structure validation for macromolecular crystallography. *Acta Crystallogr D Biol Crystallogr* **66**, 12-21 (2010). https://doi.org:10.1107/s0907444909042073
29 Lee, W., Tonelli, M. & Markley, J. L. NMRFAM-SPARKY: enhanced software for biomolecular NMR spectroscopy. *Bioinformatics* **31**, 1325-1327 (2015). https://doi.org:10.1093/bioinformatics/btu830
30 Punjani, A., Rubinstein, J. L., Fleet, D. J. & Brubaker, M. A. cryoSPARC: algorithms for rapid unsupervised cryo-EM structure determination. *Nat Methods* **14**, 290-296 (2017). https://doi.org:10.1038/nmeth.4169
31 Punjani, A., Zhang, H. & Fleet, D. J. Non-uniform refinement: adaptive regularization improves single-particle cryo-EM reconstruction. *Nat Methods* **17**, 1214-1221 (2020). https://doi.org:10.1038/s41592-020-00990-8
32 Pettersen, E. F. *et al.* UCSF Chimera--a visualization system for exploratory research and analysis. *J Comput Chem* **25**, 1605-1612 (2004). https://doi.org:10.1002/jcc.20084
33 Jo, S., Kim, T., Iyer, V. G. & Im, W. CHARMM-GUI: a web-based graphical user interface for CHARMM. *J Comput Chem* **29**, 1859-1865 (2008). https://doi.org:10.1002/jcc.20945
34 Lee, J. *et al.* CHARMM-GUI Input Generator for NAMD, GROMACS, AMBER, OpenMM, and CHARMM/OpenMM Simulations Using the CHARMM36 Additive Force Field. *J Chem Theory Comput* **12**, 405-413 (2016). https://doi.org:10.1021/acs.jctc.5b00935
35 Phillips, J. C. *et al.* Scalable molecular dynamics on CPU and GPU architectures with NAMD. *J Chem Phys* **153**, 044130 (2020). https://doi.org:10.1063/5.0014475
36 Humphrey, W., Dalke, A. & Schulten, K. VMD: visual molecular dynamics. *J Mol Graph* **14**, 33-38, 27-38 (1996). https://doi.org:10.1016/0263-7855(96)00018-5
37 Pronk, S. *et al.* GROMACS 4.5: a high-throughput and highly parallel open source molecular simulation toolkit. *Bioinformatics* **29**, 845-854 (2013). https://doi.org:10.1093/bioinformatics/btt055
38 Villinger, S. *et al.* Functional dynamics in the voltage-dependent anion channel. *Proc Natl Acad Sci U S A* **107**, 22546-22551 (2010). https://doi.org:10.1073/pnas.1012310108
39 Miehling, J., Goricanec, D. & Hagn, F. A Split-Intein-Based Method for the Efficient Production of Circularized Nanodiscs for Structural Studies of Membrane Proteins. *Chembiochem* **19**, 1927-1933 (2018). https://doi.org:10.1002/cbic.201800345
40 Waterhouse, A. M., Procter, J. B., Martin, D. M., Clamp, M. & Barton, G. J. Jalview Version 2--a multiple sequence alignment editor and analysis workbench. *Bioinformatics* **25**, 1189-1191 (2009). https://doi.org:10.1093/bioinformatics/btp033
